# MYCN mediates cysteine addiction and sensitizes to ferroptosis

**DOI:** 10.1101/2021.08.10.455675

**Authors:** Hamed Alborzinia, Andrés F. Flórez, Sina Gogolin, Lena M. Brückner, Chunxuan Shao, Moritz Gartlgruber, Michal Nadler-Holly, Matthias Ziehm, Franziska Paul, Sebastian Steinhauser, Emma Bell, Marjan Shaikhkarami, Sabine Hartlieb, Daniel Dreidax, Elisa M. Hess, Jochen Kreth, Gernot Poschet, Michael Büttner, Barbara Nicke, Carlo Stresemann, Jan H. Reiling, Matthias Fischer, Ido Amit, Matthias Selbach, Carl Herrmann, Stefan Wölfl, Kai-Oliver Henrich, Thomas Höfer, Frank Westermann

## Abstract

Aberrant expression of MYC family members predicts poor clinical outcome in many human cancers. Oncogenic MYC profoundly alters metabolism and mediates an antioxidant response to maintain redox balance. Here we show that MYC induces massive lipid peroxidation upon depletion of cysteine, the rate-limiting amino acid for glutathione biosynthesis and sensitizes cells to ferroptosis, an oxidative, non-apoptotic and iron-dependent type of cell death. In *MYCN*-amplified childhood neuroblastoma, MYCN mediates resistance to ferroptosis by activating transsulfuration of methionine to cysteine. MYCN may contribute to spontaneous tumor regression in low-risk neuroblastomas by promoting ferroptosis in cells with epigenetically silenced cystathionine-beta-synthase, the rate-limiting enzyme for transsulfuration. We identified enzymes and antiporter proteins crucial to ferroptotic escape, providing multiple previously unknown sites that may be acted on therapeutically.

## Introduction

Many human cancers depend on aberrant expression of MYC transcription factor family members for unrestricted growth and proliferation, with high expression levels predicting poor clinical outcome^1–4^. How MYC-mediated metabolic adaptations ensure nutritional uptake and synthesis of cellular building blocks required for rapidly expanding cell populations remains poorly understood. Experimental evidence^5–8^ has long been interpreted as addiction of MYC-driven cancer cells to the amino acid glutamine, which when absent causes apoptosis. The finding that MYC can promote cell death as well as cell proliferation remains paradoxical in view of MYC oncogenic potential. Transgenic mouse models support MYC requirement for both tumor development and functions triggering massive cell death depending on tissue type and context^5^. Whether these cell death-promoting activities are irreversibly impaired by mutations in MYC-driven cancer cells or may be reactivated remains to be clarified.

Childhood neuroblastoma, an embryonic tumor derived from progenitors of the sympathetic nervous system, is a paradigmatic model for MYC-driven cancers^6^. *MYCN* amplification identifies a highly aggressive subtype associated with malignant progression and poor outcome despite intensive multimodal therapy^7,8^. In contrast, a relatively large fraction of low-risk neuroblastomas with elevated MYCN expressed from a normal *MYCN* locus, particularly those arising in children under 18 months of age, regress spontaneously even if the disease is metastatic^9,10^. While the forms of regulated cell death involved are unclear, genes required for apoptosis, such as caspase 8 (*CASP8*), are often epigenetically silenced and anti-apoptotic signals, such as expression of the *MDM2* proto-oncogene, are active in primary high-risk and relapse neuroblastomas^11–13^.

## Results

### Cystine deprivation induces ferroptosis

We analyzed the interplay of oncogenic MYCN or MYC, referred to here as MYC(N), activity with amino acid metabolism. Proliferation was slowed without inducing death (Extended Data Fig. 1a, b) by downregulating MYCN to a ‘MYCN-low’ state (~65% reduction, Fig. 1a) in the *MYCN*-amplified IMR5/75 neuroblastoma cell model^14^, and intracellular pools of all amino acids were reduced (Fig. 1b), most prominently cysteine up to tenfold (Fig. 1c). Inhibiting MYCN binding to MYC associated factor X (MAX) with 10058-F4^15^ achieved similar results (Fig. 1b and Extended Data Fig. 1c). These experiments suggest that an oncogenic ‘MYCN-high’ cellular background heavily utilizes cysteine. Systematically depleting individual amino acids from the growth medium confirmed the selective dependence of cells in the ‘MYCN-high’ state on cystine (oxidized form of cysteine imported by cells and readily reduced to two cysteine molecules). Cystine deprivation caused cell death preferentially in ‘MYCN-high’ cells, and was more detrimental in this context than the previously described glutamine deprivation^16–19^ (Fig. 1d). Instituting a ‘MYCN-low’ cellular background inhibited cell death induced by cystine deprivation (Fig.1d). Moreover, inhibiting MYCN-MAX binding for MYCN function largely rescued cystine-deprived ‘MYCN-high’ cells from cell death (Fig. 1e). Overexpressing MYCN in a diploid *MYCN* background using the Tet21N neuroblastoma cell model^20^ (Fig. 1a) further confirmed cystine addiction in cells with a ‘MYCN-high’ state (Fig. 1f). We stably integrated a regulable element capable of reducing MYC protein levels by ~85% (Fig. 1a) in the *MYC*-amplified NCI-H23 lung cancer cell line, and showed that cystine deprivation also caused cell death in the ‘MYC-high’ cellular background that could be largely prevented by MYC knockdown (Fig. 1g). These data suggest that survival of MYC(N)-driven tumor cells acutely depends on cystine availability, which maintains an adequate intracellular cysteine supply.

**Figure 1.**
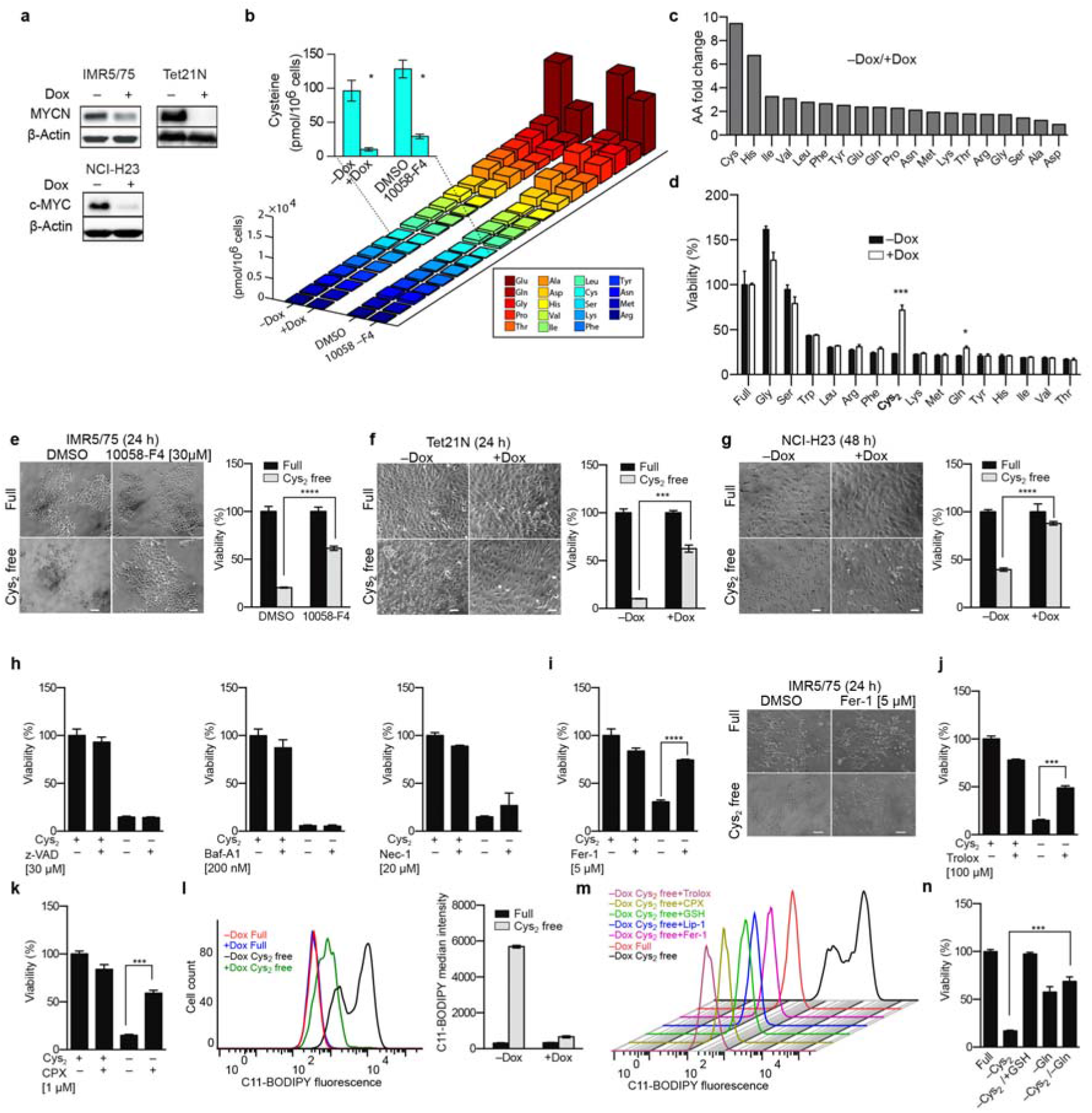
Cystine addiction of MYC(N)-expressing neuroblastoma and lung cancer cells. **a**, Representative western blot of neuroblastoma (IMR5/75, Tet21N) and lung cancer (NCI-H23) cells upon MYC(N) knockdown using doxycycline (– Dox, high expression; +Dox, low expression). **b, c**, Intracellular amino acid quantification (**b**) and ranked fold changes (**c**) in IMR5/75 reveal cysteine (Cys) reduction after MYCN inhibition (+Dox or 10058-F4, 96 h). **d**, Standardized viability of IMR5/75 after single amino acid depletions (48 h) reflects cell death resistance of cystine (Cys_2_)-deprived ‘MYCN-low’ cells. **e-g,** Cys_2_ addiction is linked to ‘MYC(N)-high’ state; Scale: 50 μm, mean of triplicates standardized to NT (full medium). **h-k**, Cys_2_ deprivation-induced death of IMR5/75 is preventable using ferroptosis inhibitors. **l-n**, Flow cytometry confirms L-ROS accumulation in Cys_2_-deprived ‘MYCN-high’ IMR5/75 (**l**), reduction by ferroptosis inhibitor treatments or GSH supplementation (**m**) and maintained viability (**n**) when supplied with GSH or glutamine (Gln, 24 h). Data are representative of at least two (**e-g**), three (**d, h-n**) or more (**b**) biological replicates (means ± SD). two-tailed Student’s t-test: *P□<□0.05, **P□<□0.01, ***P□<□0.001, ****P□<□0.0001.

To investigate the form of regulated cell death induced by cystine deprivation, we used small molecule inhibitors. Inhibiting caspase proteases to prevent apoptosis, lysosomal function to inhibit autophagy, or RIPK1 to prevent necroptosis, did not abolish IMR5/75 cell death in the ‘MYCN-high’, cystine-deprived state (Fig. 1h). Cystine uptake is required for the synthesis of glutathione, a major antioxidant preventing elevated levels of reactive oxygen species (ROS). An oxidative, non-apoptotic, and iron-dependent form of regulated cell death caused by ROS-mediated massive lipid peroxidation (L-ROS) was recently described called ferroptosis^21,22^. Death of neuroblastoma cells with the oncogenic ‘MYCN-high’ background in a cystine-deprived state was averted by a specific inhibitor of ferroptosis or the lipophilic antioxidant, trolox (Fig. 1i, j). Supplementing cystine-free medium with the intracellular iron chelator, ciclopirox olamine (CPX), also prevented death of neuroblastoma cells with the ‘MYCN-high’ cellular background (Fig. 1k). Thus, intracellular cysteine depletion in a ‘MYCN-high’ context induces neuroblastoma cell death by ferroptosis.

To investigate L-ROS formation in the context of cystine deprivation, we stained IMR5/75 cells with the lipid peroxidation sensor, C11-BODIPY, then flow cytometrically analyzed L-ROS in either ‘MYCN-high’ or ‘MYCN-low’ backgrounds with or without access to cystine from the growth medium. Cellular L-ROS was not altered by MYCN expression levels alone, but was approximately tenfold higher in the ‘MYCN-high’ than ‘MYCN-low’ background following cystine deprivation (Fig. 1l). Inhibiting ferroptosis with either ferrostatin-1 or liproxstatin-1 in the ‘MYCN-high’, cystine-deprived state rescued cells from massive L-ROS accumulation, as did treatment with the lipophilic antioxidant, trolox, or supplementing cystine-free medium with the intracellular iron chelator, CPX (Fig. 1m). Supplementing cystine-free medium with glutathione also averted L-ROS accumulation and cell death (Fig. 1m), pinpointing glutathione as a crucial factor preventing L-ROS accumulation and ferroptosis in the oncogenic ‘MYCN-high’ and cysteine-deprived context. Paradoxically, depletion of both glutamine and cystine delayed ferroptosis in the ‘MYCN-high’ state (Fig. 1n). It is widely thought that glutamine deprivation drives MYC-dependent cells into apoptosis by creating an intracellular glutamate deficiency^16–19^. Our data only show a slight selective effect of glutamine deprivation on cell death in the ‘MYCN-high’ state and no additive effect for depletion of glutamine and cystine. New data gathered in mouse embryonic fibroblasts show that glutaminolysis is essential for ferroptosis triggered by depletion of full amino acids or cystine alone^23^. Our findings together with new descriptions of the ferroptotic process establish a novel functional link between oncogenic MYC(N) and ferroptosis, and imply regulation by cysteine-dependent glutathione availability.

### Glutathione becomes an Achilles’ heel

Cells in the ‘MYCN-high’ state may be susceptible to redox imbalances and respond by maintaining high glutathione levels to prevent iron-dependent L-ROS formation and subsequent ferroptosis. MYC has been reported to upregulate glutathione in other cancer cells^24^. Neuroblastoma cells in the ‘MYCN-high’ state had three-to fourfold higher intracellular glutathione levels (Fig. 2a) and twofold higher reduced:oxidized glutathione ratios (Fig. 2b), reducing intracellular ROS levels (Fig. 2c) compared with the ‘MYCN-low’ state. An unbiased high-throughput MYCN synthetic lethal siRNA screen (Fig. 2d) identified genes preferentially acting in the ‘MYCN-high’ state and protecting cells from ROS accumulation and ferroptosis. Top hits included six enzymes in glutathione metabolism, which are preferentially active in lipid peroxide detoxification (Fig. 2e–f and Extended Data Table 1). Massive reduction in cell number was selectively triggered in the ‘MYCN-high’ context (Fig. 2f, g) by individual knockdown of glutathione-disulfide reductase (*GSR*), glutathione peroxidase 4 or 6 (*GPX4*, *GPX6*), or glutathione S-transferase mu 1, mu 5, or kappa 1 (*GSTM1*, *GSTM5*, *GSTK1*). Knockdown of genes for the two enzymes catalyzing glutathione biosynthesis, glutamate-cysteine ligase catalytic subunit (*GCLC*) and glutathione synthetase (*GSS*) were synthetic lethal with MYCN, but were not prioritized under the stringent criteria applied in our unsupervised statistical selection (Fig. 2h). Glutathione peroxidase (*GPX4*) was among our top hits, and is a potent negative regulator of ferroptosis^22^. Co-treatment with the MYC-MAX inhibitor, 10058-F4, rescued cells in the ‘MYCN-high’ state from undergoing ferroptosis in response to RSL-3, a small molecule inhibiting GPX4 (Fig. 2i). Five neuroblastoma cell lines harboring various *MYCN* complements and expressing MYC(N) protein levels ranging from non-detectable to strong responded differently to cystine deprivation (Extended Data Fig. 2a, b). MYC(N) expression in these cell lines determined their response to RSL-3 treatment, with the SK-N-FI cell line, lacking detectable MYC(N) expression, being virtually resistant to ferroptosis induction (Fig. 2j). Elevated cellular free iron levels have been suggested to predispose cancer cells to ferroptosis^21,22^, and oncogenic MYC induces iron uptake required for diverse enzymatic reactions in fast-proliferating cells by activating the transferrin receptor (*TFRC)^25^*. We hypothesized that increasing the iron supply may boost ferroptotic cell death in MYCN-driven cells. Indeed, supplying iron enhanced the RSL-3 effect on the *MYCN*-amplified KELLY cell line by ~tenfold (Fig. 2k). Taken together, we identify multiple points in glutathione synthesis and metabolism, particularly detoxification of L-ROS, that are vulnerable in the ‘MYCN-high’ context, and show that ferroptosis is dependent on MYC(N) expression and enhanced by iron.

**Figure 2.**
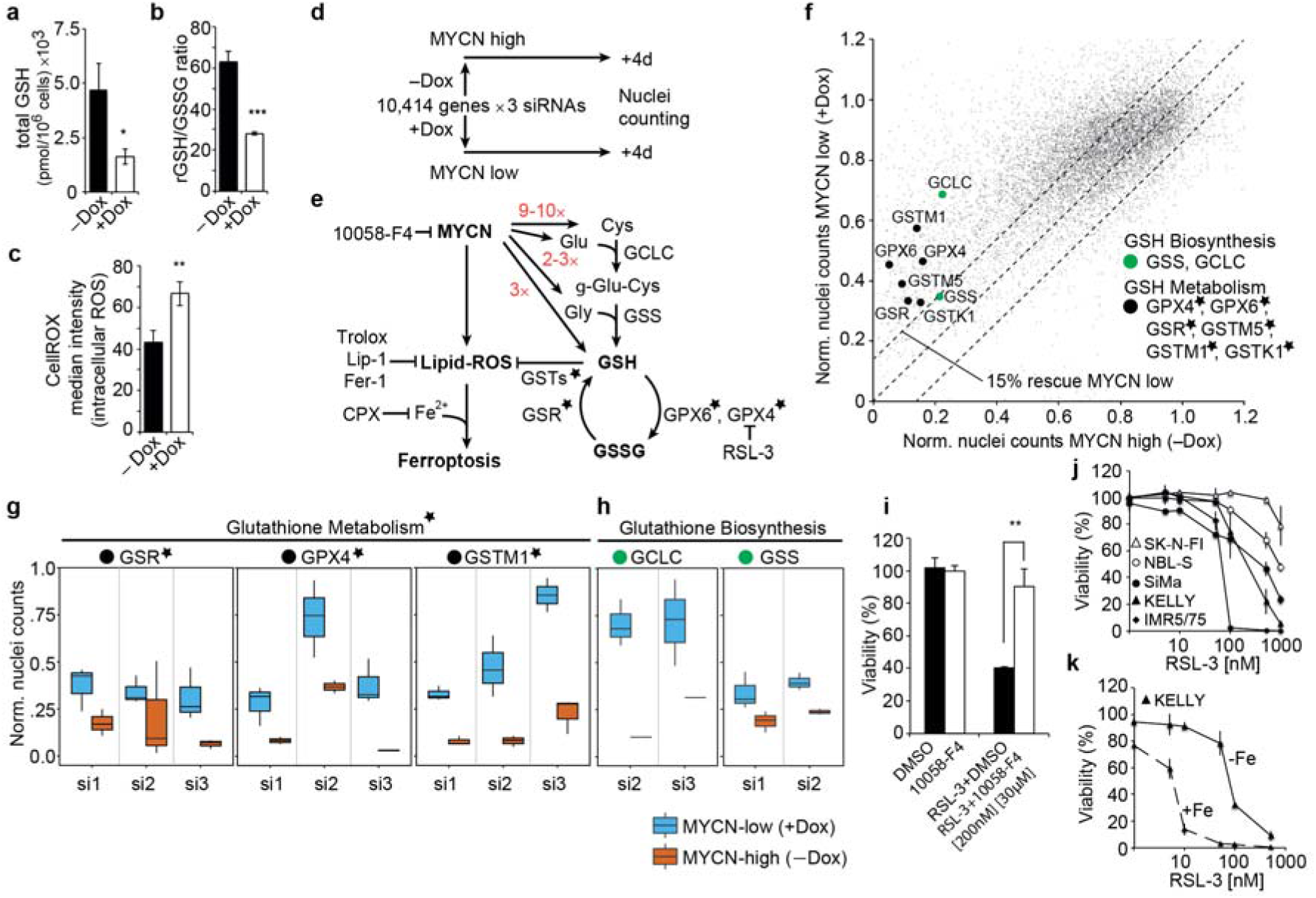
Inhibition of GSH metabolism is synthetic lethal with high MYCN. **a**, HPLC reveals elevated intracellular glutathione, and **b**, ratio of reduced:oxidized GSH expressed as GSH equivalents in ‘MYCN-high’ IMR5/75. **c**, Flow cytometry confirms intracellular ROS reduction (lower CellROX intensity) in ‘MYCN-high’ state; three biological replicates; means ± SD. **d-h**, MYCN synthetic lethal druggable genome-wide siRNA screening approach in IMR5/75 (**d**); MYCN effects on L-ROS formation and intracellular amino acid levels (fold changes in red), the two-step biosynthesis of GSH and GSH metabolism; *top MYCN synthetic lethal hits of GSH metabolism, false discovery rate (FDR) of 0.2; action of ferroptosis inhibitors (CPX, Fer-1, Lip, Trolox and 10058-F4) and class II ferroptosis inducer (RSL-3) as indicated (**e**); **f**, Effects of individual siRNAs (grey dots): ‘MYCN-high’ vs. ‘MYCN-low’, including key players (median of 2-3 siRNAs) of **g**, top MYCN synthetic lethal hits ( *) of GSH metabolism (black) and **h**, GSH biosynthesis (green). **i-k**, 10058-F4 treatment prevents RSL-3-induced cell death in IMR5/75 (72 h) (**i**); MYCN expression level of neuroblastoma cell lines reflects capacity to resist RSL-3-induced ferroptosis: cells with *MYCN*-amplification (black symbols), moderate MYCN expression, (white circles) and lack thereof (white triangles) (**j**), Fe supplementation (25 μg/ml) enhances RSL-3-induced death in KELLY (**k**). n = 3 (four technical replicates); \**P*□<□0.05 \*\**P*□<□0.01. \*\*\**P*□<□0.001; means ± SD; two-tailed Student’s t-test.

### Transsulfuration maintains glutathione

Cellular cysteine can be oxidized from imported cystine or synthesized from methionine^26^. The solute carrier family 7 member 11 (SLC7A11) of the x_c_^-^ system imports cystine in exchange for glutamate^27^, whereas SLC7A5 imports methionine and branched chain amino acids in exchange of glutamine^28^ (Fig. 3a). Sulfasalazine and erastin are small molecule inhibitors of SLC7A11, from which erastin also inhibits SLC7A5^28–30^. Selective induction of ferroptosis in ‘MYCN-high’ cells, compared to cells in the ‘MYCN-low’ state or with MYC(N) activity inhibited by 10058-F4, returned with erastin treatment (Fig. 3b and Extended Data Fig. 3a). Erastin also induced death in a similar pattern to RSL-3 in the five neuroblastoma cell lines expressing a panorama of oncogenic MYCN levels (Fig. 3b) that was boosted by iron supply (Extended Data Fig. 3b), demonstrating that erastin induces ferroptosis in the ‘MYCN-high’ cellular context. SLC7A5 knockdown revealed a MYCN synthetic lethal effect, whereas SLC7A11 knockdown had a similar effect in both ‘MYCN-high’ and ‘MYCN-low’ contexts (Fig. 3c), as did treatment with sulfasalazine (Extended Data Fig. 3c). We conclude that quantitatively reducing SLC7A11 antiporter activity does not limit cysteine availability. MYC(N)-selective induction of ferroptosis appears to be independent of cystine uptake by SLC7A11 and conferred by the import of methionine in exchange for glutamine by SLC7A5.

**Figure 3.**
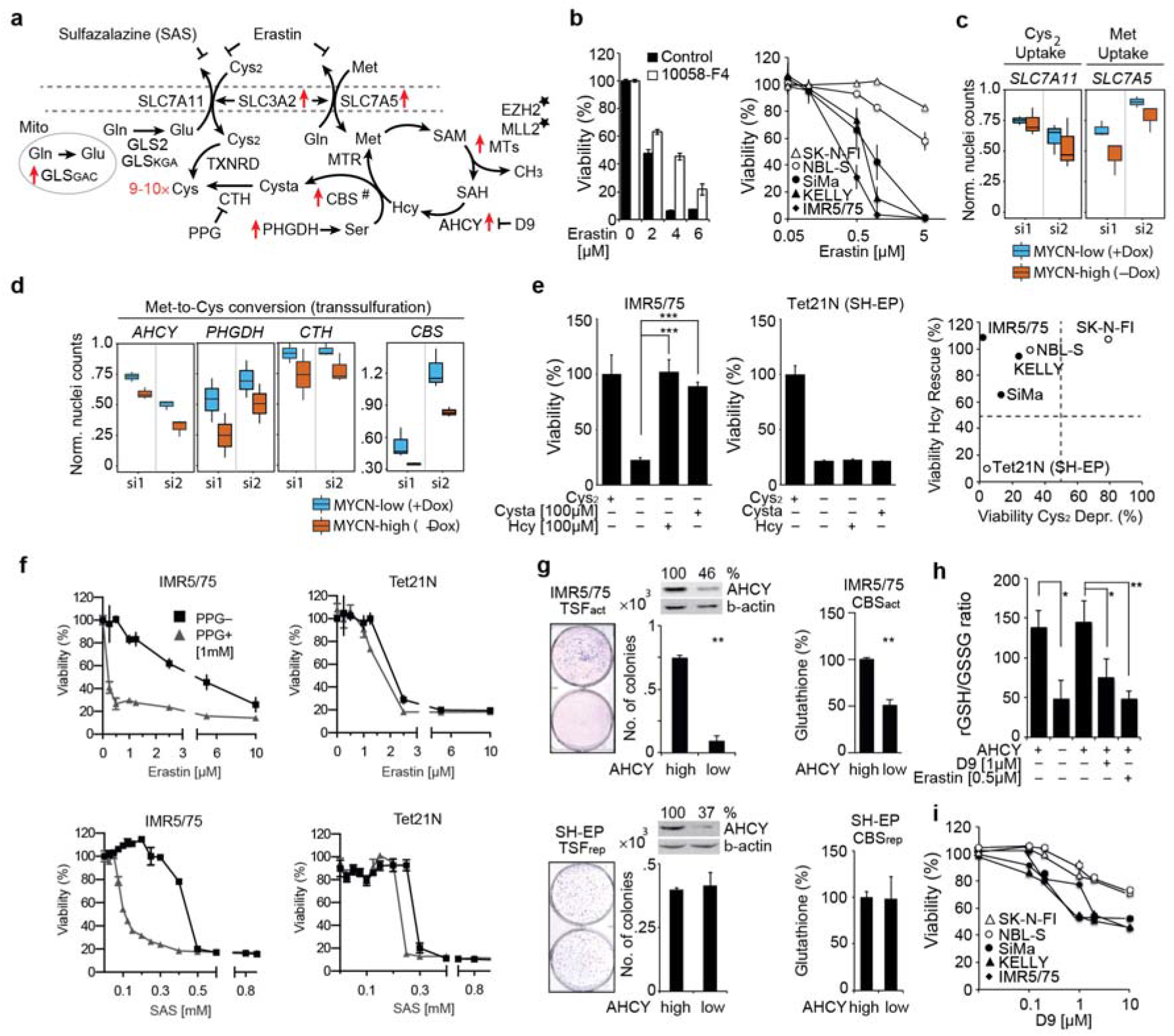
MYCN activates transsulfuration genes controlling Met-to-Cys conversion in *MYCN*-amplified cells. **a**, Illustration of Cys_2_ and Met uptake/metabolism, and transsulfuration (^#^), Mito = mitochondria, MTs = methyltransferases, Cysta = cystathioinine; *: top hits from MYCN siRNA screen; red arrows: genes transcriptionally activated by MYCN in IMR5/75 (FDR 0.001, using likelihood ratio testing) with Cys fold changes (red). **b**, Erastin treatment (72 h) enables cell death in ‘MYCN-high’ IMR5/75, which is inhibited by 100058-F4, and proves more potent in neuroblastoma cell lines with high MYCN activity; n = 3 (four technical replicates). **c**, **d**, IMR5/75 MYCN siRNA screen shows MYCN-independent *SLC7A11* knockdown effect but MYCN synthetic lethality with knockdown of *SLC7A5* (**c**) and of key transsulfuration genes (**d**); two individual siRNAs. **e**, Homocysteine (Hcy) and Cysta supply prevent Cys_2_ deprivation-induced ferroptosis in ‘MYCN-high’ IMR5/75 and cell lines with naturally evolved MYCN dependency; n = 3 (four technical replicates). **f**, Inhibition of Cys synthesis (PPG) enables erastin- or SAS-induced cell death in MYCN-dependent IMR5/75, unlike in ectopically MYCN expressing TET21N, n=3, representative experiment is shown. **g**-**i**, AHCY inhibition (knockdown or D9) reduces colony formation, glutathione synthesis (**g**) and, like erastin, the rGSH/GSSG ratio in IMR5/75 (**h**). D9 is more potent in cell lines with high MYC(N) expression (**i**); n = 3; \**P*□<□0.05 \*\**P*□<□0.01 \*\*\**P* <0.001; means ± SD; two-tailed Student’s t-test.

Cysteine is synthesized from methionine via transsulfuration, whereby homocysteine, a methionine cycle intermediate, and serine are combined to cystathionine and then converted to cysteine^26^ (Fig. 3a). Four genes encoding enzymes key for converting methionine to cysteine were synthetic lethal with MYCN (Fig. 3d), as were two methyltransferases that may indirectly increase homocysteine levels (Extended Data Fig. 3c). We interpret these findings to mean that transsulfuration provides a critical additional cysteine source for *MYCN*-amplified cells. Supporting this interpretation, supplementing media with either the homocysteine or cystathionine intermediate prevented cystine deprivation-induced ferroptosis in IMR5/75 cells in the ‘MYCN-high’ state (Fig. 3e) and cell lines maintaining oncogenic MYCN activity (Fig. 3e, Extended Data Fig. 4). Activating transsulfuration appears to require cell type-specific factors present in cells that have naturally evolved dependency on oncogenic MYCN signaling, since supplying homocysteine or cystathionine to a cell model expressing the *MYCN* transgene in the normal diploid *MYCN* background did not avert cystine deprivation-induced ferroptosis (Fig. 3e). Pharmacologically inhibiting intracellular cysteine synthesis with an inhibitor of cystathionine-gamma-lyase (PPG) sensitized cells with naturally evolved MYCN dependency and active transsulfuration, but not ectopically expressed MYCN and inactive transsulfuration, to erastin- or sulfasalazine-induced cell death (Fig. 3f). Downregulating S-adenosylhomocysteine hydrolase (*AHCY*) reduced glutathione levels and ratios of reduced:oxidized glutathione states (Fig. 3g, h). Colony formation by cells in the ‘MYCN-high’ state was impaired relative to the amount of *AHCY* downregulation (Fig. 3g). In contrast, *AHCY* downregulation had no effect on glutathione levels or colony formation (Fig. 3g) in cells with a diploid *MYCN* background and inactive transsulfuration. Similar to erastin and RSL-3, pharmacologically suppressing homocysteine synthesis by inhibiting AHCY with D9^31^ in five neuroblastoma cell lines induced cell death in a manner positively correlated with MYCN expression level (Fig. 3i). Thus, neuroblastoma cells with oncogenic *MYCN* activity use the methionine cycle and transsulfuration as an internal cysteine source for glutathione biosynthesis to evade ferroptosis.

### Oncogenic MYCN induces transsulfuration

Our data show that both cystine import and intracellular cysteine synthesis achieve the intracellular state supportive of oncogenic MYC(N)-driven growth without endangering the cell to ferroptosis. To understand how cysteine metabolism and redox homeostasis are adapted by oncogenic MYC(N) activity, we analyzed differential gene expression in synchronized cells in ‘MYCN-high’ and ‘MYCN-low’ states^32^. Synchronizing the cell cycle separated indirect effects changing with cell cycle phases and related to accelerated proliferation from oncogenic MYC(N) actions pervasive throughout the cell cycle (Fig. 4a, Extended Data Fig. 5a). Expression of cystathionine-beta-synthase (*CBS*), *AHCY* and phosphoglycerate dehydrogenase (*PHGDH*), three key enzymes for transsulfuration, was elevated in the ‘MYCN-high’ state across all cell cycle phases, while expression of the cysteine uptake-controlling x_c_^-^ system (*SLC7A11*) was unaffected by MYCN expression level (Fig. 4a and Extended Data Fig. 5a). Control by oncogenic MYC(N) of the three transsulfuration enzymes was confirmed in a panel of 29 neuroblastoma cell lines with diverse *MYC(N*) expression (Extended Data Fig. 5b). Histone marks associated with active transcription depict MYCN as an amplifier of active *AHCY* transcription in cells with oncogenic MYC(N) activity (Extended Data Fig. 6a, b). *CBS* regulation appears from ChIP-seq analysis to be orchestrated by both activating and silencing histone modifications. The *CBS* regulatory region was bound by MYCN and harbored activating histone marks, while almost completely lacking silencing marks, in cells with oncogenic *MYCN* (Fig. 4b, c, Extended Data Fig. 7a, b). The *CBS* promoter was predominantly occupied by repressive marks in cells harboring diploid *MYCN* and lacking detectable MYC(N) expression (Fig. 4b, c, Extended Data Fig. 7a, b). Ectopic MYCN expression in the *MYCN* diploid background induced expression of *AHCY* and other MYCN targets, while *CBS* remained epigenetically repressed (Extended Data Fig. 7c). Differences in *CBS* expression in primary neuroblastomas correlated with histone modifications and methylation of intronic CpGs dependent on the genomic *MYCN* status (Fig. 4d, Extended Data Fig. 8). These data paint a picture where *AHCY* is constitutively active in neuroblastoma cells, but further enhanced by MYCN, while *CBS* is more strictly epigenetically controlled and may require prior release from epigenetic repression before MYCN-dependent activation can occur.

**Figure 4.**
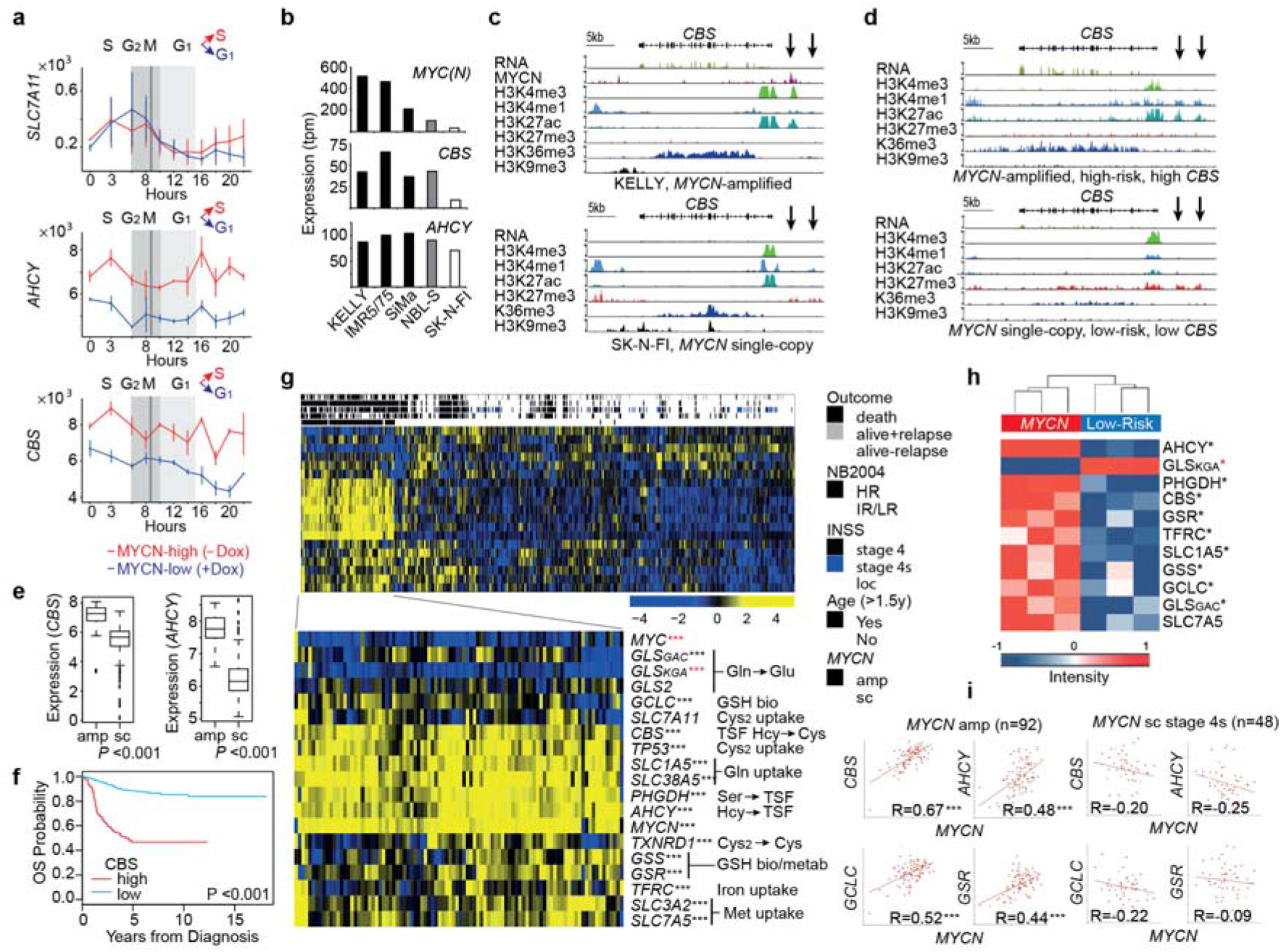
Differential anti-ferroptotic activities in regressing and progressing neuroblastomas. **a**, Expression of transsulfuration genes is linked to MYCN expression level during cell cycle progression, unlike SLC7A11 expression (IMR5/75, RNA-seq); means ± SD; differentially expressed genes were identified using likelihood ratio testing. **b**, mRNA expression of cells reflecting diverse MYC(N) expression levels. **c**, **d**, mRNA expression (RNA-seq) and input normalized read counts of histone modifications and MYCN (ChiP-seq) at the *CBS* locus in representative *MYCN*-amplified and *MYCN* single-copy cells (**c**) and tumors (**d**). **e**, *MYCN* status-dependent expression in 498 primary neuroblastomas (92 *MYCN*-amplified, 406 *MYCN* single-copy tumors, RNA-seq), Wilcoxon rank-sum test. **f**, Kaplan-Meier overall survival estimates for subgroups defined by *CBS* expression, high *CBS* (n=123); low *CBS* (n=375). **g**, Gene expression heatmap of subgroups hierarchically clustered with average linkage and noncentralized correlation distance function; row: transcript, column: sample, HR = high-risk, IR/LR = intermediate/low-risk according the German NB2004 study, amp = amplified, sc = single-copy, *** (black) *P* <0.001 higher and *** (red) *P* <0.001 lower gene expression, Wilcoxon rank-sum test. **h**, Mass spectrometry reveals differential expression of proteins mediating GSH synthesis/metabolism and GSH-relevant amino acid uptake/metabolism: *MYCN*-amplified vs. low-risk *MYCN* single-copy tumors, * (black) *P* <0.05 higher protein, * (red) *P* <0.05 lower protein expression using t-test, intensities as z-scores. **i**, Pearson’s correlation analysis of *MYCN* expression and GSH synthesis/metabolism and transsulfuration genes in *MYCN*-amplified (n=92) and *MYCN* single-copy stage 4s (n=48) tumors.

Global gene expression profiles from 498 primary neuroblastomas^33^ also showed significantly higher *CBS* and *AHCY* expression in *MYCN*-amplified neuroblastomas (Fig. 4e, g). *MYCN* expression strongly correlated with *CBS* and *AHCY* expression in tumors, and high *CBS* or *AHCY* levels were associated with poorer overall patient (Fig. 4f, g and Extended Data Figure 9). Mass spectrometry-based global proteomes from six primary neuroblastomas also showed that CBS, AHCY and PHGDH expression was higher in tumors harboring *MYCN* amplifications (Fig. 4h). *CBS* and *AHCY* expression correlated with *PHGDH* expression in tumors as well as with *TP53* expression, which is only mutated in 2% of primary neuroblastomas^34^ (Fig. 4g). *TP53* is a transcriptional target of MYCN^10,35^ and is known to suppress cystine uptake via the x_c_^-^ system by *SLC7A11* repression^36^. *SLC7A11* expression did not correlate with either *MYCN* amplification, *TP53* expression or other risk factors for poor patient outcome (Fig. 4g). Combining these data with the unchanged expression of the x_c_^-^ system in cells in either the ‘MYCN-high’ or ‘MYCN-low’ states, confirms a lack of direct control of cystine uptake by oncogenic MYCN activity, but may indicate that processes simultaneously up- and downregulating *SLC7A11* may be holding expression steady in a varying MYC(N) context. Although the mechanisms upregulating the enzymes for transsulfuration in a cell with oncogenically active MYC(N) may require indirect components, MYC(N) drives increased transsulfuration activity, rather than cysteine import, in tumor cells to maintain the cellular cysteine supply for glutathione synthesis.

The ‘MYCN-high’ state also supported enhanced methionine, glutamine and iron uptake throughout the cell cycle (Extended Data Fig. 5a). Expression of transsulfuration enzymes in primary neuroblastomas also correlated with transporters for glutamine, methionine, and iron uptake (Fig. 4g). Enhanced expression of transporter proteins for methionine, glutamine, and iron was confirmed in *MYCN*-amplified tumor proteomes (Fig. 5h). Because genes mediating cytosolic glutaminolysis^37^ were barely expressed in neuroblastoma cells and tumors (Extended Data Fig. 5c, Fig. 4g), it is unlikely that imported glutamine is used as a source for glutamate in cysteine exchange via the x_c_^-^ system and glutathione synthesis occurring in the cytosol^24^. Higher expression of the mitochondrial glutaminolysis isoform^38^ in *MYCN*-amplified tumor transcriptomes and proteomes (Fig. 4g, h) suggests that glutamate synthesis is required only in the neuroblastoma cell mitochondria. Expression of two enzymes regulating the glutathione biosynthesis (*GCLC*) and redox cycle (*GSR*) positively correlated with MYC(N) expression in neuroblastoma cell lines (Extended Data Fig. 5b) and were more strongly expressed in the *MYCN*-amplified tumor transcriptomes and proteomes (Fig. 4g, h). Oncogenic MYC(N) activity appears to fuel hyperactive glutathione redox activity by increasing import of methionine, upregulating three key enzymes for transsulfuration to increase intracellular cysteine and activating glutathione biosynthesis and metabolism.

**Figure 5.**
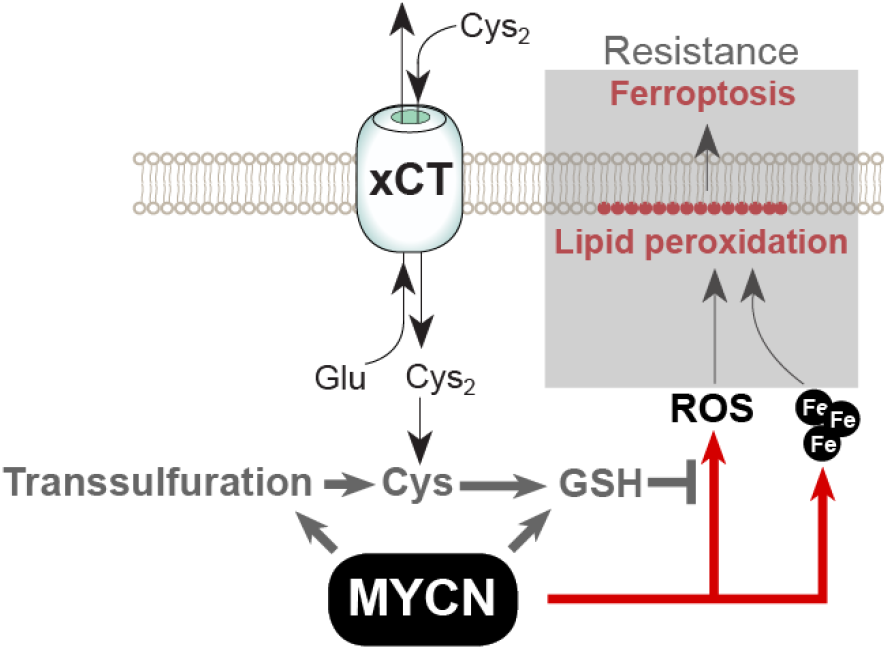
MYC(N)-mediated ferroptosis resistance. Cells with oncogenic MYC(N) respond to reduction of cysteine levels with massive lipid peroxidation resulting in ferroptosis, an oxidative, non-apoptotic and iron-dependent type of cell death. Deregulation of key components acting in transsulfuration, glutathione redox activity (grey arrows) and methionine import provides MYCN-driven neuroblastoma cells and tumors essential escape mechanisms to ferroptosis.

Elevated MYCN expression from a normal *MYCN* locus occurs in low-risk neuroblastomas, particularly in metastatic disease that tends to regress spontaneously in children under 18 months of age^10^. Glutathione biosynthesis and redox activity, which climbs in the *MYCN*-amplified scenario, remained unchanged with fluctuating MYCN in low-risk, metastatic neuroblastomas (Fig. 4i). Moreover, intronic CpG methylation epigenetically silenced *CBS* in low-risk, metastatic cases with low *CBS* expression (Extended Data Fig. 8), indicating that transsulfuration is not available to increase internal cysteine for glutathione biosynthesis.

## Discussion

We show that MYCN sensitizes neuroblastoma cells to an oxidative, non-apoptotic, iron-dependent regulated type of cell death when intracellular cysteine availability for glutathione synthesis is limited. We propose that a ‘MYCN-high’ state in neuroblastoma cells sensitizes them to lipid peroxidation, which in combination with an ‘acute’ intracellular cysteine reduction, triggers metabolic sabotage^39^ and massive ferroptotic cell death that may explain spontaneous neuroblastoma regression (Fig. 5). Robust molecular markers for tissues undergoing ferroptosis are not yet known^40^. However, our study shows transsulfuration, glutathione redox activity and methionine import to be crucial to escape ferroptosis, and correlated expression of key genes for these processes with MYCN expression in tumors. Expression for these key genes did not correlate with MYCN expression in low-risk primary neuroblastomas that frequently undergo spontaneous regression, and repressive histone marks at *CBS* regulatory regions and hypermethylated intronic CpGs were associated with low *CBS* expression in the low-risk neuroblastomas. The absence of MYCN-dependent epigenetic regulation of *CBS* in a low-risk neuroblastoma compounded by reduced import of methionine would prevent the tumor from developing the hyperactive glutathione biosynthesis and redox activity it needs to guard against ferroptosis. Spontaneous regression may be the physiological resolution of this cellular state. Unlike low-risk tumors, *MYCN*-amplified neuroblastomas appear to metabolically adapt to survive events resulting in intracellular cysteine reduction, such as TP53 activation and SLC7A11 repression^36^ following telomere crisis. The cysteine requirement of cancers dependent on oncogenic MYC(N) activity creates a previously unknown Achilles’ heal that could be exploited to selectively induce ferroptosis for treatment. Our findings identify cysteine uptake, intracellular methionine conversion to cysteine, the L-ROS-specific scavenging system, and the cellular free iron pool as vulnerabilities in cancer cells driven by oncogenic MYC(N) activity such as *MYCN*-amplified neuroblastomas.

## Extended data figures

**Extended data figure 1.**
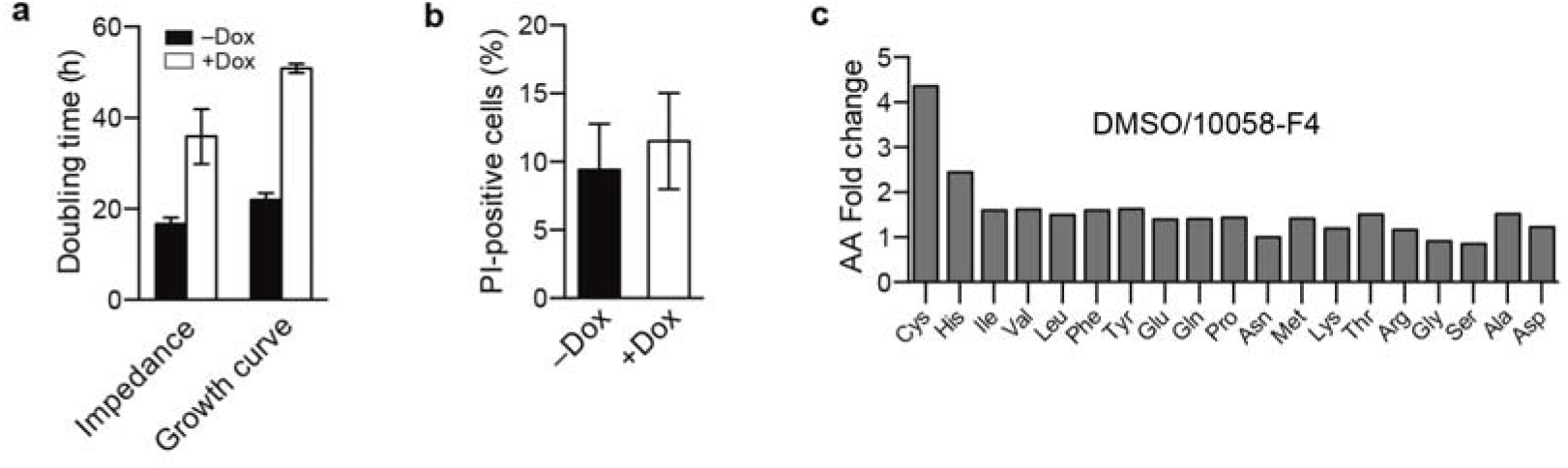
**a**, Doubling time calculation from exponential growth curve quantified by FACS and impedance using the RCTA system for ‘MYCN-high’ (–Dox) and ‘MYCN-low’ (+Dox) IMR5/75 cells. Cell proliferation curves of exponentially growing cells (two biological replicates; means ± SD). **b**, Viability quantification using FACS and propidium iodide staining in exponentially growing cells: – Dox vs. +Dox. **c**, Fold changes of intracellular amino acid levels 24 h after 10058-F4 treatment (normalized with DMSO control).

**Extended data figure 2.**
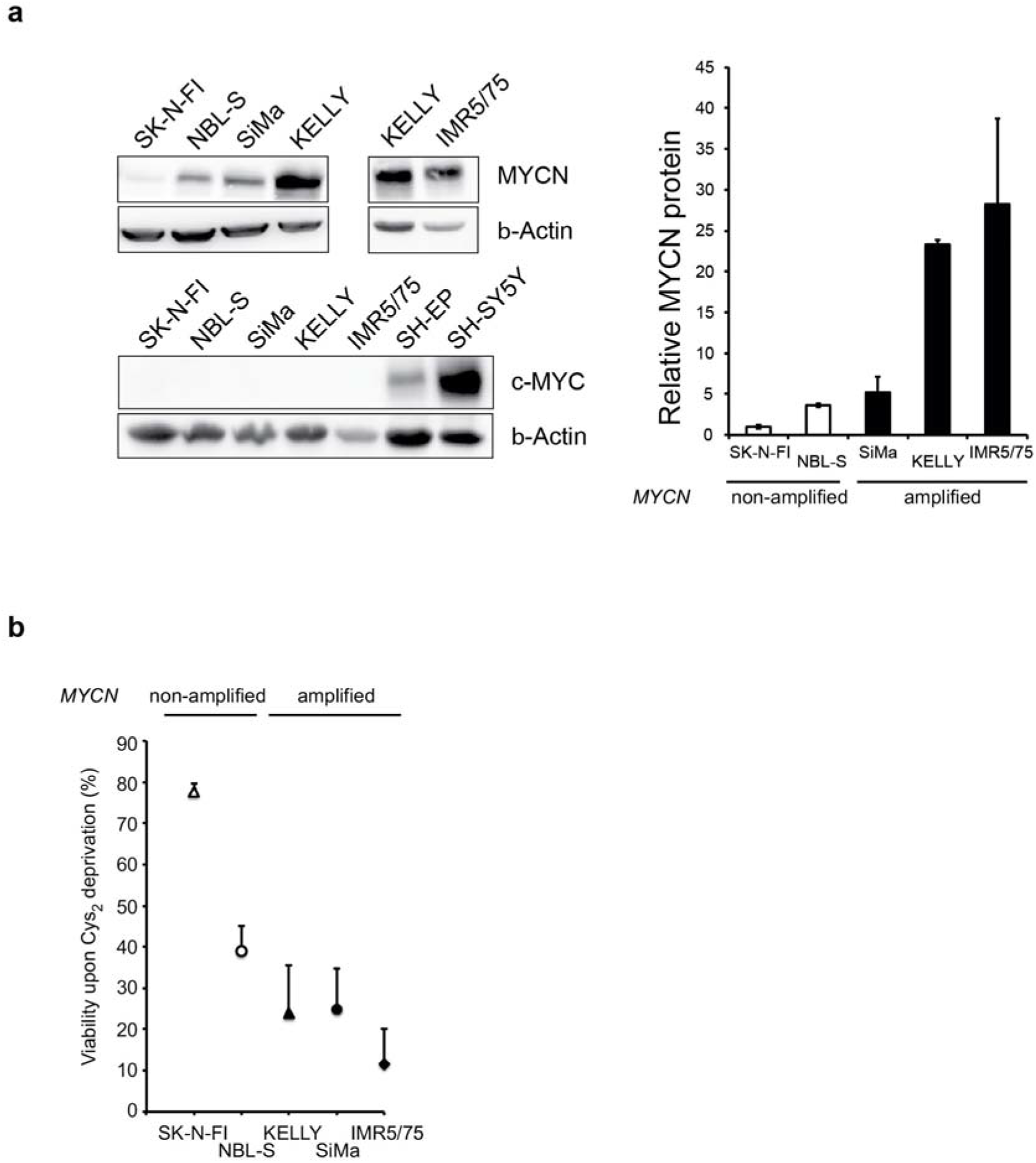
**a**, MYCN and c-MYC protein expression in five representative neuroblastoma cell lines with diverse MYC(N) expression levels (western blot). □-actin was used as loading control. SH-SY5Y and SH-EP cells were used as positive control for high and low c-MYC-expressing neuroblastoma cell lines, respectively. Quantification of MYCN protein levels; the lowest MYCN/□-actin ratio in SK-N-FI was set to one and the higher MYCN/□-actin ratios of the other cell lines calculated as fold changes. Means and standard deviations based on three independent western blot experiments. **b**, Relative viability changes (%) after 48 h Cys_2_ deprivation (% CTB fluorescence reads at 540/580 nm) for five neuroblastoma cell lines with diverse MYC(N) expression levels. Mean and standard deviation of three independent experiments are depicted. *MYCN*-amplified cell lines (black symbols), *MYCN* single-copy NBL-S cells with moderate MYCN expression (white circles), and SK-N-FI cells lacking MYC(N) expression (white triangles); n = 3 (4 technical replicates).

**Extended data figure 3.**
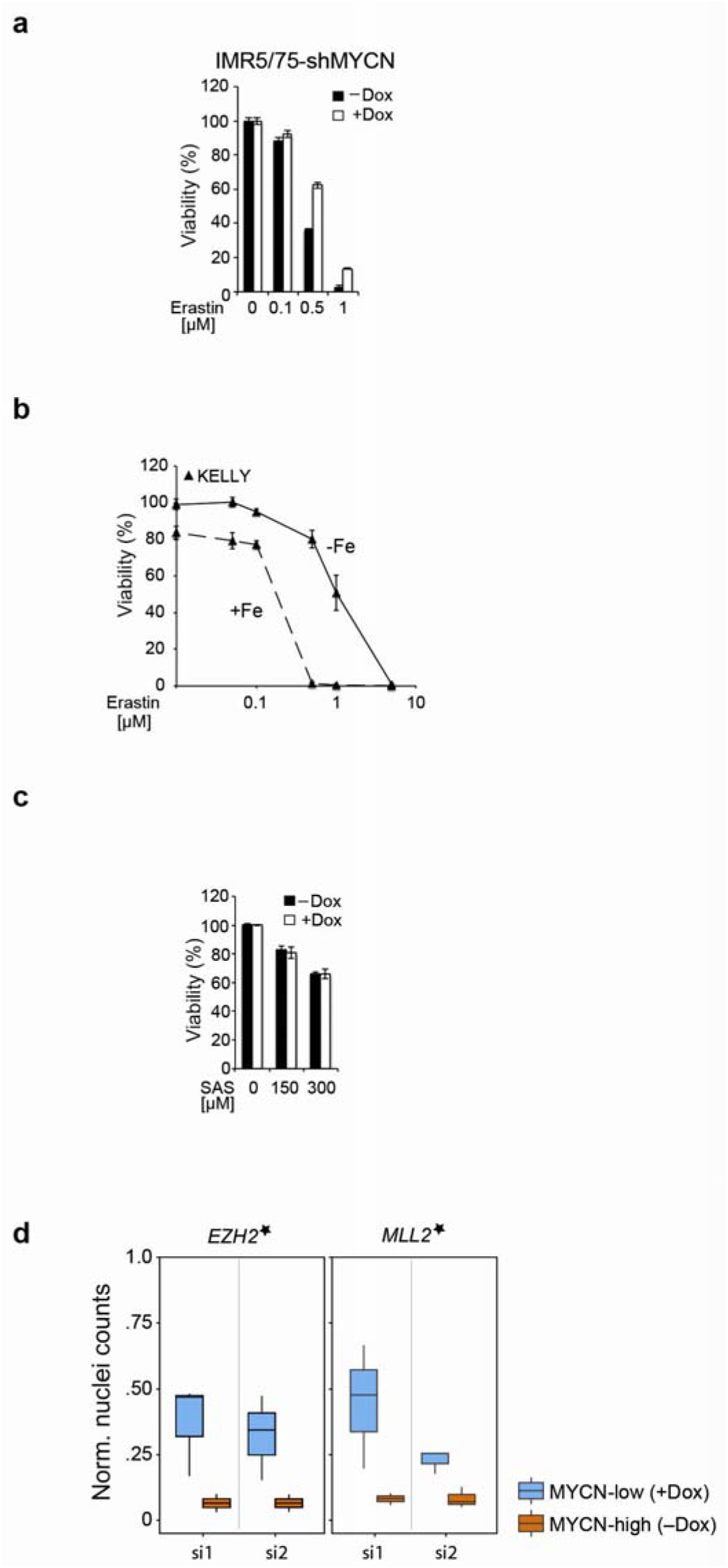
**a**, MYCN synthetic lethal effect of class I ferroptosis inducer (FIN) erastin on viability of *MYCN*-amplified IMR5/75 cells with Dox-inducible expression of a MYCN shRNA; n = 3 (4 technical replicates; means ± SD). **b**, Viability of KELLY cells upon treatment with erastin w/o iron (25 μg/ml, Venofer); n = 3 (4 technical replicates; means ± SD). **c**, Viability of IMR5/75 cells upon treatment with sulfusalazine (SAS); n = 3 (4 technical replicates; means ± SD). **d**, Two methyltransferases that may indirectly increase homocysteine levels are synthetic lethal with high MYCN (from the MYCN siRNA screen; *FDR of 0.2).

**Extended data figure 4.**
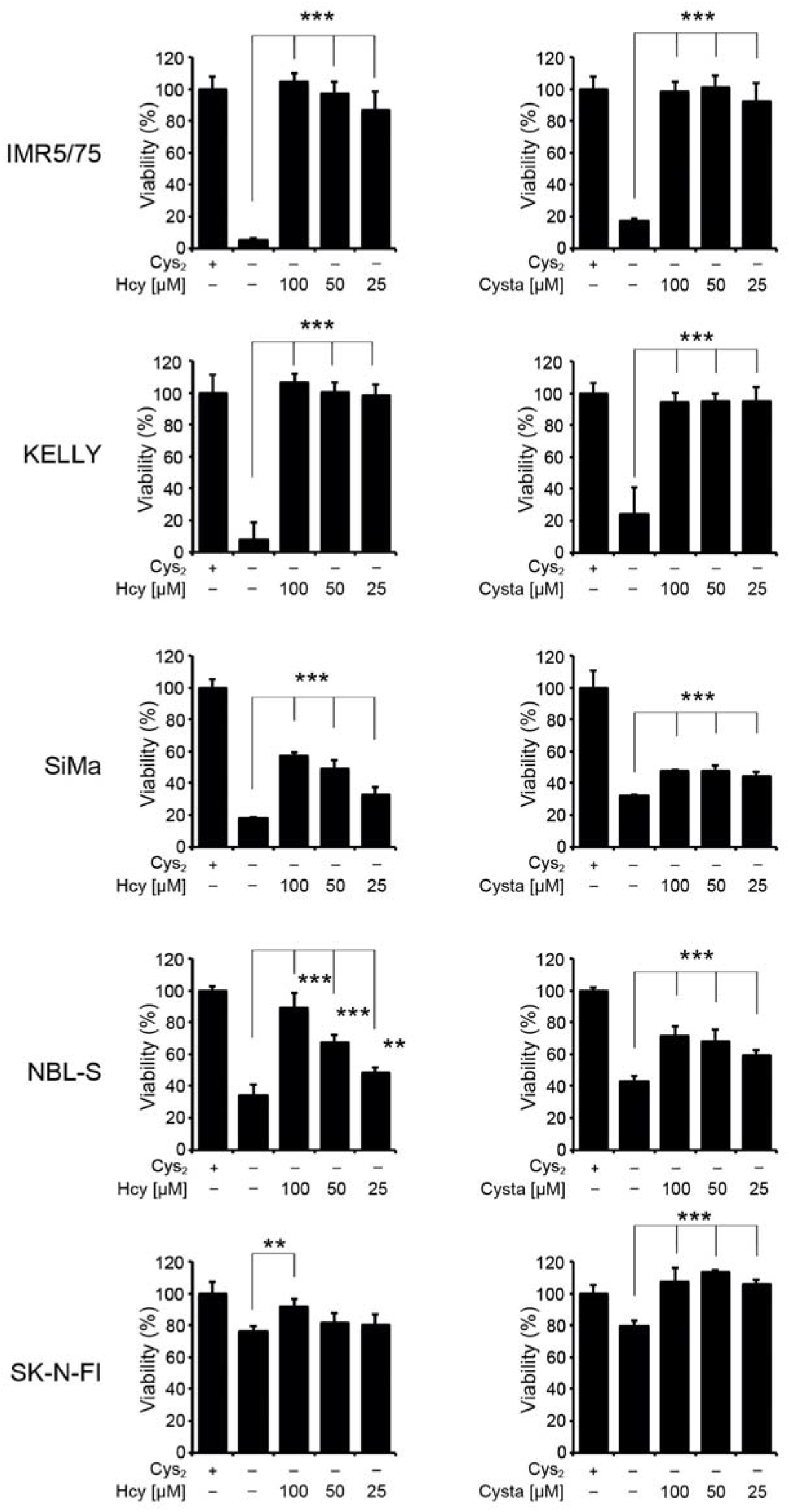
Viability response of Cys_2_-deprived neuroblastoma cell lines with different levels of MYC(N) expression to Hcy (left) or Cysta (right) treatment at indicated concentrations (vs. +Cys_2_ control); n = 3 (4 technical replicates; means ± SD); \*\**P*□<□0.01, \*\*\**P*□<□0.001; two-tailed Student’s **t**-test.

**Extended data figure 5.**
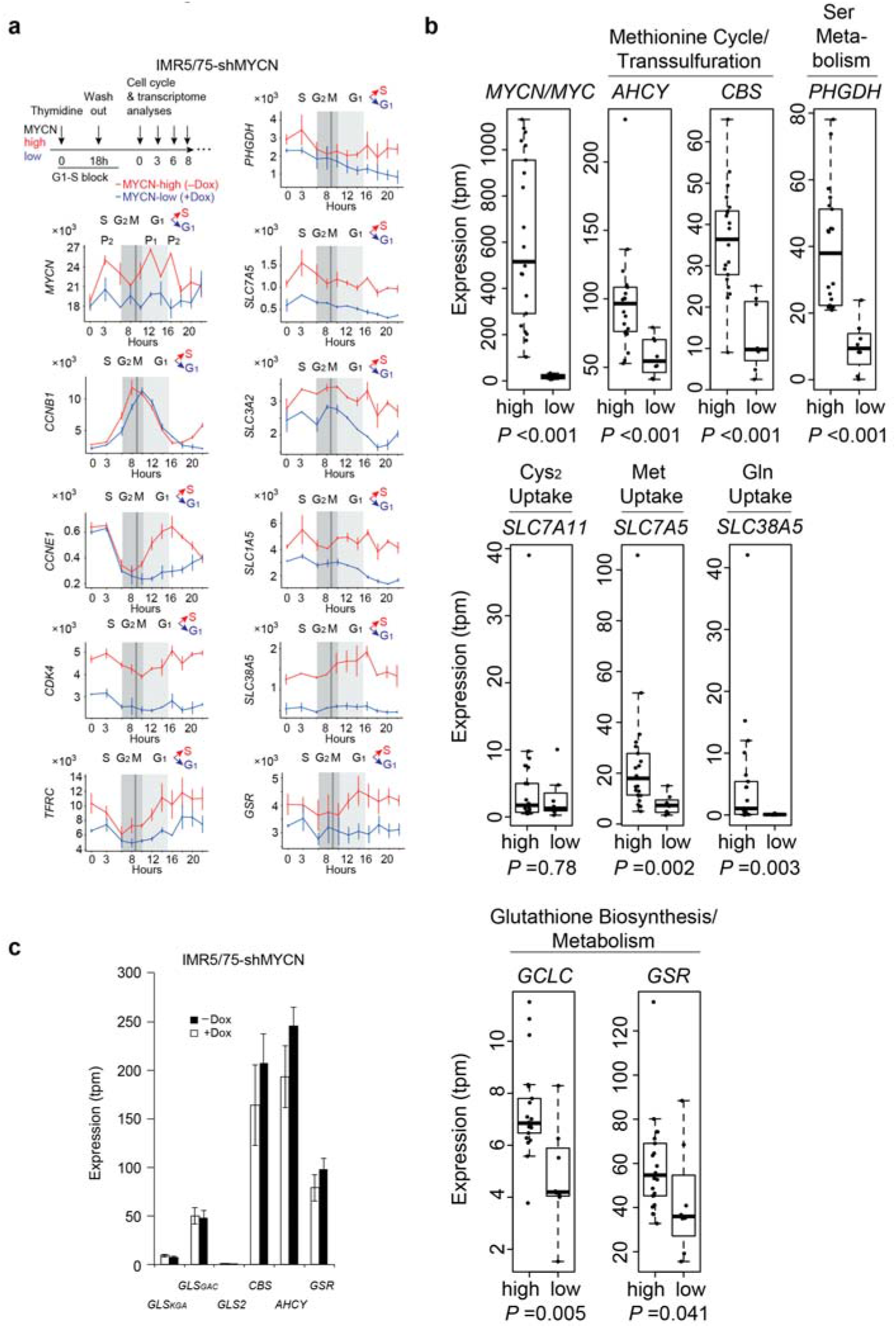
**a**, Time-resolved gene expression profiles during cell cycle progression: MYCN-high vs. MYCN-low IMR5/75 cells. To distinguish pervasive MYCN functions from indirect effects related to accelerated cell proliferation, ‘MYCN-high’ and ‘MYCN-low’ cells were synchronized using a thymidine block for 18 h. Thymidine block-released cells were harvested every 2 h over a period of 22 h for transcript profiling using RNA-seq and cell cycle analysis (flow cytometry). The *MYCN* gene expression profile reveals two cell cycle-related peaks (P1 before G1-S transition and P2 before S-G2/M). Characteristic profiles for a pervasively MYCN-induced gene (*CDK4*) and two MYCN-induced cell cycle-regulated genes (*CCNE1* for G1-S transition and *CCNB1* for G2-M transition); means ± SD. **b,** Expression of GSH biosynthesis/metabolism, transsulfuration, and amino acid uptake/metabolism genes in 29 neuroblastoma cell lines as defined by RNA-seq: MYC(N)-expressors vs. MYC(N)-non-expressors (c-MYC- and MYCN-expressing cell lines combined), Wilcoxon rank-sum test. **c**, Expression of glutaminolysis genes (mitochondrial *GLS_GAC_*; cytosolic *GLS_KGA_* and *GLS2*) as compared to MYCN-regulated genes (*CBS*, *AHCY*, *GSR*) in ‘MYCN-high’ and ‘MYCN-low’ IMR5/75 cells.

**Extended data figure 6.**
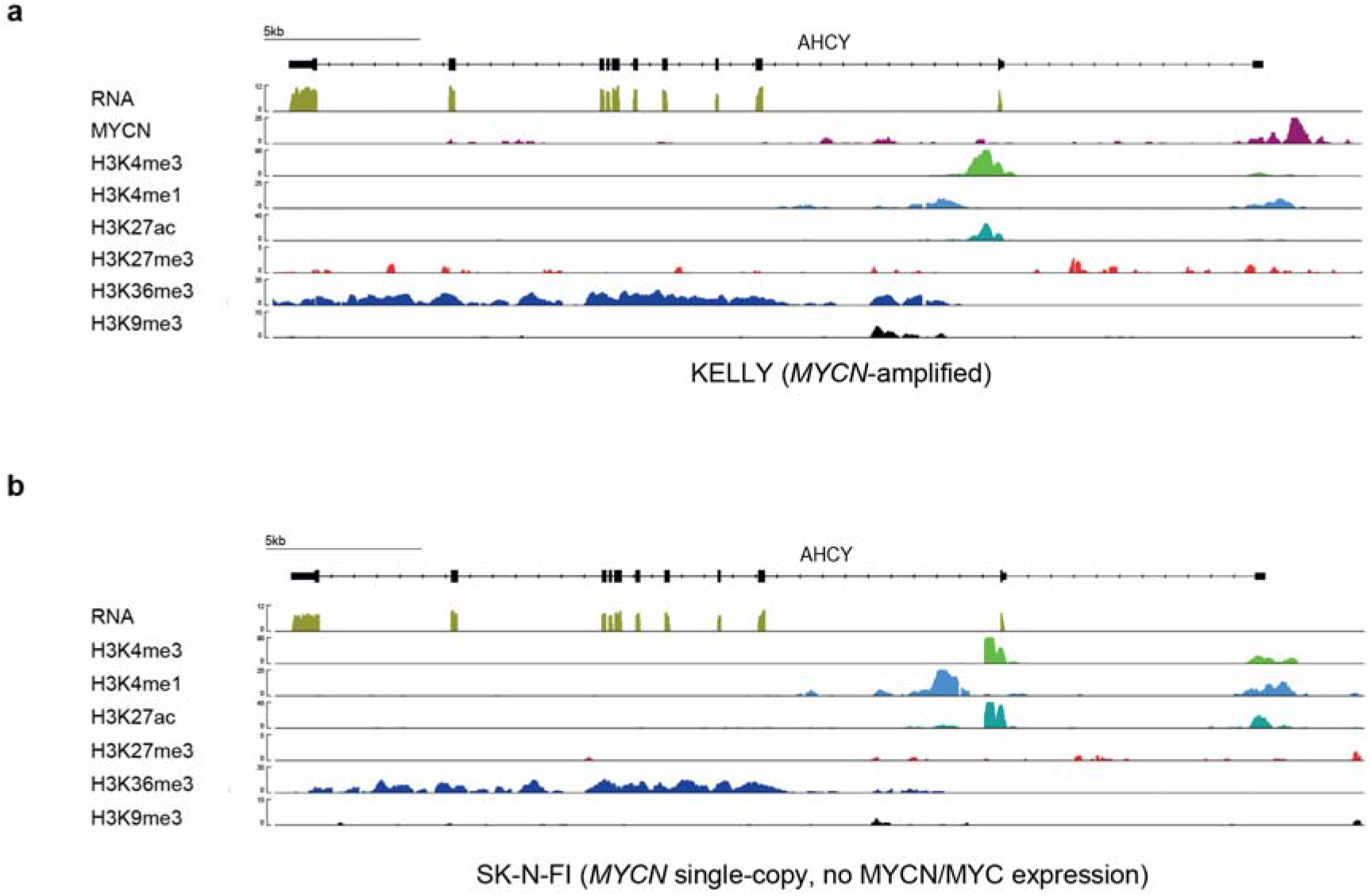
**a, b,** RNA-seq normalized reads and input normalized read counts of ChIP-seq experiments for histone modifications and MYCN at the *AHCY* gene locus in *MYCN*-amplified KELLY cells (**a**) and in *MYCN* single-copy SK-N-FI cells (**b**) lacking MYC(N) expression.

**Extended data figure 7.**
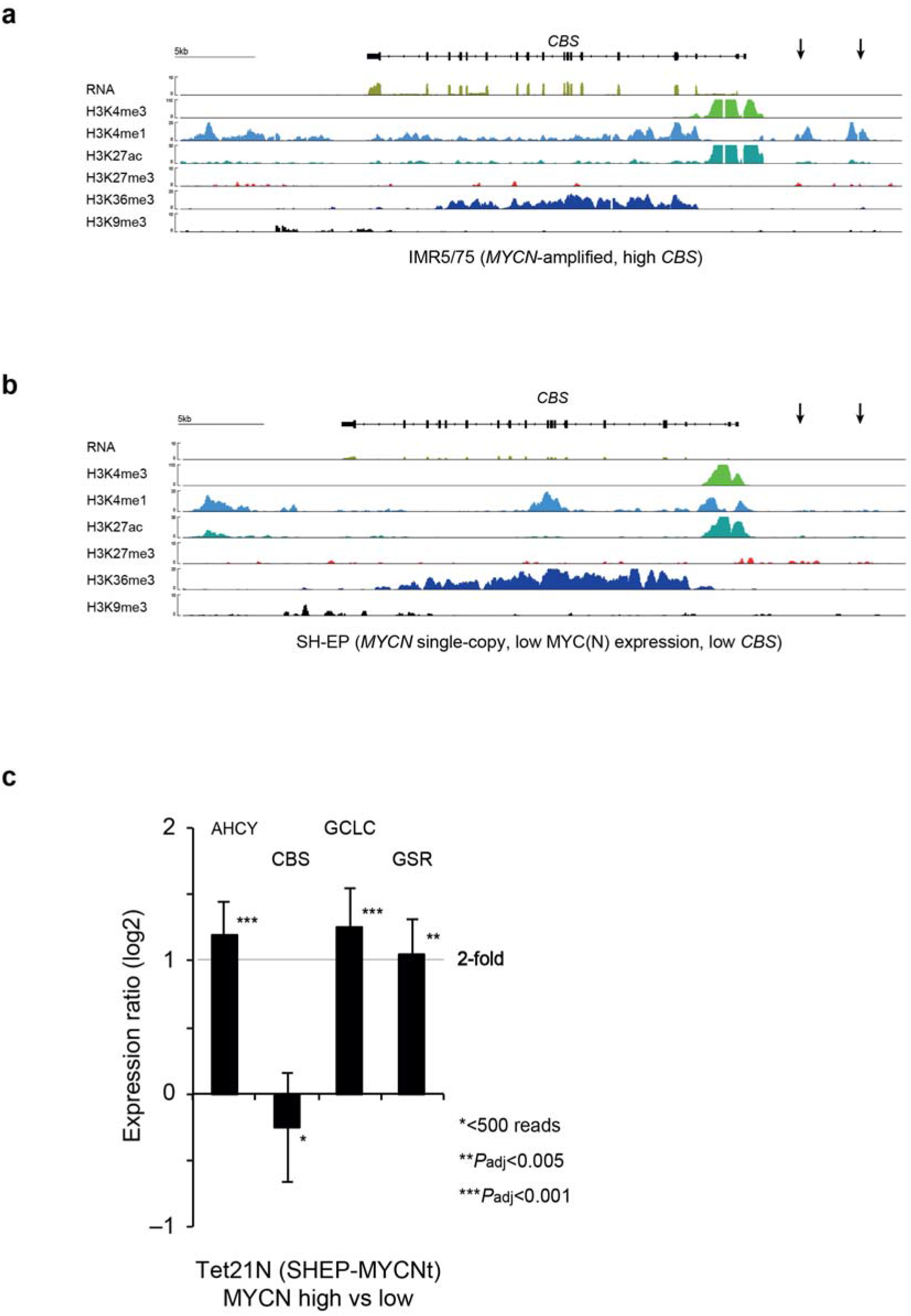
**a, b**, RNA-seq normalized reads and input normalized read counts of ChIP-seq experiments for histone modifications at the *CBS* gene locus in *MYCN*-amplified IMR5/75 cells (**a**) with high *CBS* expression and active transsulfuration and *MYCN* single-copy SH-EP cells (**b**) with low *CBS* expression and inactive transsulfuration. **c,** relative mRNA expression in Tet21N (SH-EP-MYCNt) cells harboring an inducible MYCN transgene upon MYCN induction. Tet21N and parental SH-EP cells have low *CBS* expression and inactive transsulfuration. MYCN induction has a significant effect on *AHCY*, *GCLC* and *GSR* but not on *CBS* expression; means ± SD.

**Extended data figure 8.**
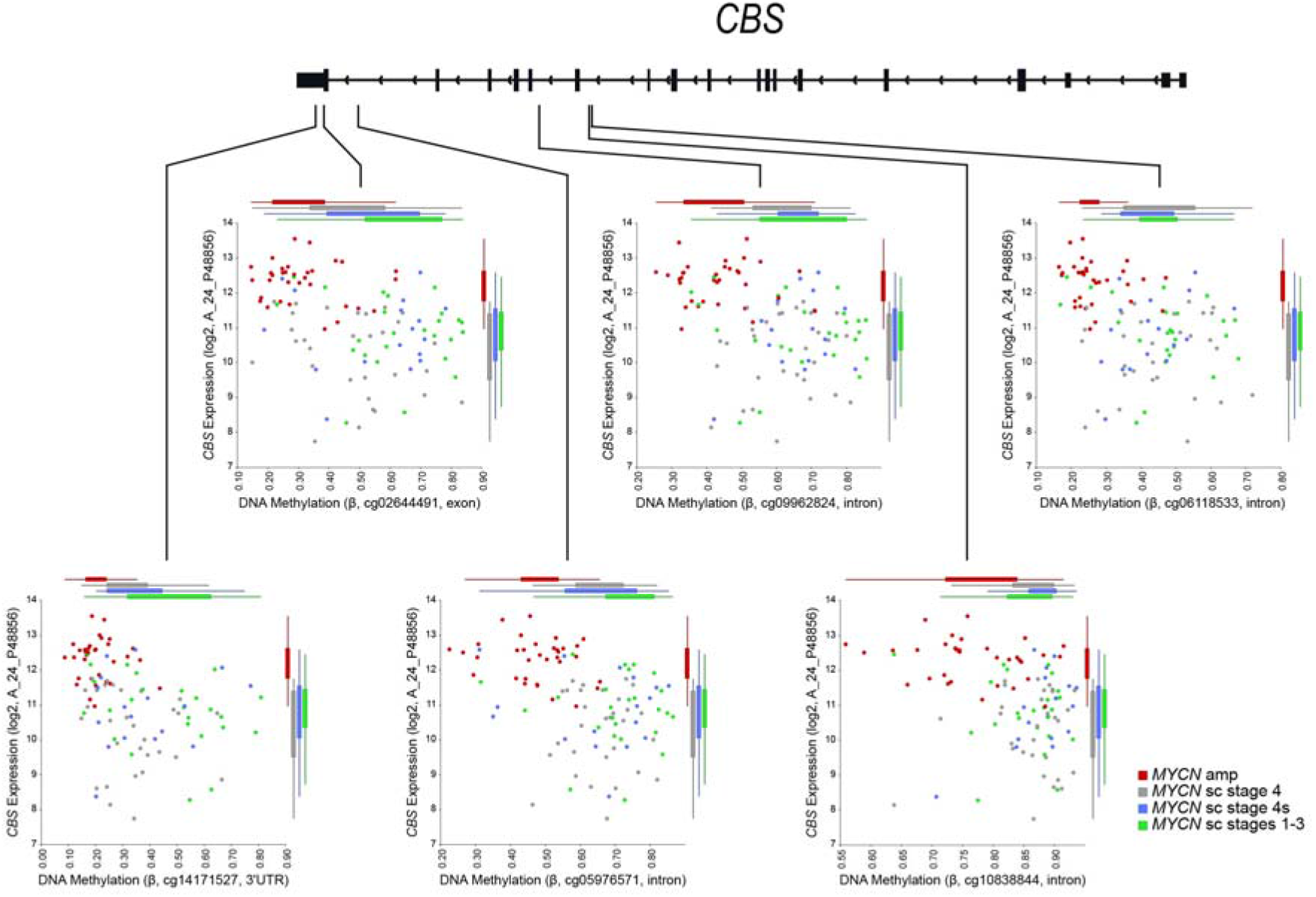
Genomic position of *CBS*-annotated CpGs whose methylation is significantly associated with *CBS* expression and patient risk (p<0.01, Wilcoxon rank-sum statistics and p<0.05, Fisher exact test). DNA methylation assessed by Infinium HumanMethylation450 BeadChips and *CBS* expression assessed by 44k customized Agilent oligonucleotide microarrays in 105 primary neuroblastomas (GEO accession GSE73518, Henrich et al. 2016) are depicted. For all *CBS* CpGs whose methylation significantly correlated with *CBS* expression and patient risk, hypomethylation was associated with *CBS* upregulation and high-risk disease. R2 Genomics Analysis and Visualization Platform (http://r2.amc.nl) was used for data visualization.

**Extended data figure 9.**
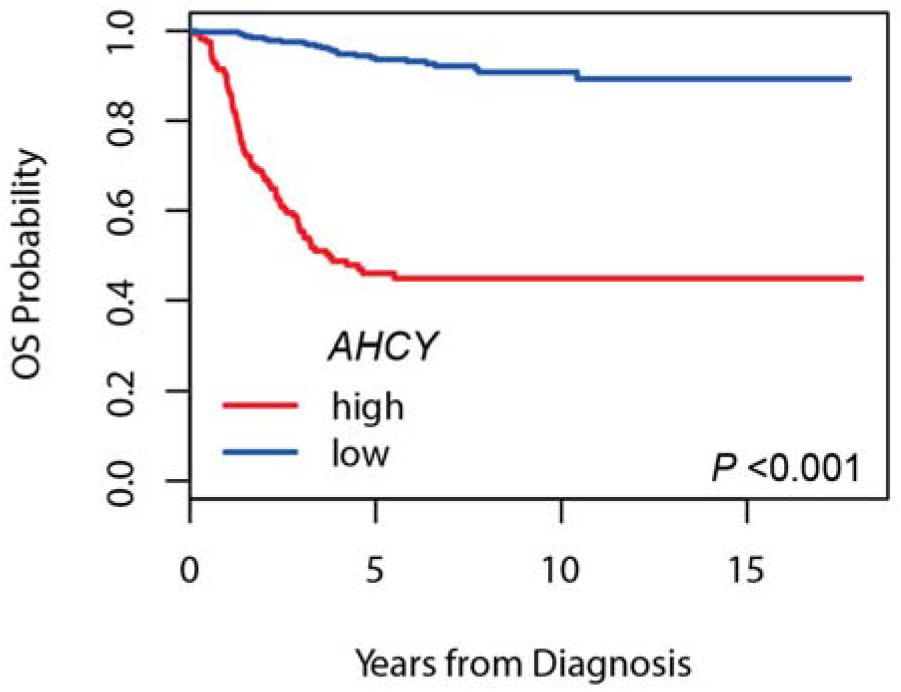
Kaplan-Meier overall survival estimates for subgroups defined by *AHCY* expression. The cutoff values for dichotomization of *AHCY* expression was estimated by maximally selected log-rank statistics, high *AHCY* expression (n=165), low *AHCY* expression (n=333).

## Acknowledgements

This work was supported by the DKFZ-BayerHealthCare Alliance (L.M.B., D.B., S.G., E.M.H. and F.W.), the e:Med initiative (SYSMED-NB, grant no. 01ZX1307D to F.W., T.H. and M. S.), the German Cancer Consortium (DKTK) Joint Funding program, the BMBF MYC-NET grant no. 0316076A to F.W. and T.H., the ERACoSysMed grant Optimize-NB to T.H. and F.W., the European Union grant no. 259348 to F.W., the German Cancer Research Center (DKFZ) intramural program for interaction projects and the DKFZ-Heidelberg Center for Personalized Oncology (HIPO) & National Center for Tumor Diseases (NCT) Precision Oncology Program (F.W.) and the Berlin Institute of Health (M. S.). S.W. was funded by BfR FKZ1329-468. M.G. and F.P. were supported by PhD fellowships of the German-Israeli Helmholtz Research School in Cancer Biology. The group of J. R. is supported by Merck KGaA. A.F.F thanks Julia Lochead and Tatjana Ryl for experimental support and Carina Thomé for help with RTCA measurements. We are indebted to the patients and their parents of making available the tumor specimens that were analyzed within this study, and we thank the German neuroblastoma biobank for providing patient samples. The Institutional Review Board (IRB) approved collection and use of all specimens in this study. We would like to thank Frank-Detlef Scholle and Sebastian Räse (Bayer AG) for automatic microscopic image analysis (MYCN SL screen), our colleagues Young-Gyu Park and Denise Brünig for technical assistance, and Kathy Astrahantseff, Tobias Dick, Maria Llamazares and Theodor C. H. Cole for critical reading of the manuscript. We thank the Metabolomics Core Technology Platform at the University of Heidelberg and Rüdiger Hell for providing data of amino acids and other metabolites. We would also like to acknowledge the DKFZ core facility, helping us to obtain high-quality data of sequencing and DNA methylation analyses.

## Contributions

The study was conceived by A.F.F, H.A and F.W. Metabolic assays and viability studies were conducted and analyzed by H.A., A.F.F., L.M.B., M.S., E.M.H., J.K. and S.G. S.G. performed the MYCN-synthetic lethal siRNA screen. M.G. helped with MYCN-synthetic lethal screen optimization. C.S., S.S., A.F.F and C.H. assisted with bioinformatic analyses. M.G., F.P., S.H., and D.D. performed and analyzed ChIP-seq and ChIPmentation experiments. M.N.-H., M.Z. and M.S. provided tumor proteomic data. E.B., M.G., E.M.H. and J.K. conducted RNA-seq experiments. Regulable cell models were established and colony formation assays performed by J.K. Western blotting was conducted by S.G., E.M.H. and J.K. G.P. and M.B. implemented and analyzed LC-MS measurements. K.-O.H. contributed DNA methylation analyses. B.N., C.S., J.H.R., M.F., I.A., S.W. contributed reagents, materials and analysis tools. The manuscript was prepared by F.W., T.H., H.A., A.F.F., L.M.B. and S.G. All authors approved the current version of the manuscript.

## Material and Methods

### Experimental in vitro procedures

#### Cell Culture

Human neuroblastoma (IMR5/75, KELLY, SiMa, NBL-S, SK-N-FI) and NCI-H23 cells (human NSCLC) were cultivated at 37°C with 5 % CO_2_ in RPMI 1640 medium (Gibco, Thermo Fisher Scientific, California, USA) supplemented with 10 % fetal calf serum (FCS; Gibco), and penicillin/streptomycin (AppliChem, Darmstadt, Germany). Tunable cell lines, IMR5/75 *MYCN* shRNA, SH-EP *MYCN* transgene (Tet21N) and NCI-H23 *MYC* shRNA were generated and cultured as described previously^14,20^. Cell line identity/unique SNP profiles were confirmed by the Multiplexion Multiplex Cell Authentication service (Heidelberg, Germany) as described recently^41^. To assess the effects of amino acid deprivation, cells were cultivated using modified amino acid-free DMEM powder (PAN-Biotech, Aidenbach, Germany) supplemented with individual amino acids (Sigma-Aldrich, Munich, Germany) as indicated, at final concentrations used in standard DMEM.

#### Analysis of Cell Viability and Proliferation

The impact of various treatments on cellular proliferation/viability was assessed using an SRB assay as described previously^42^. To determine changes in cellular proliferation, approximately 2×10^4^ cells were seeded per well (48-well format) in full medium. After 24 h, cells were washed with PBS, fed with the chosen medium and treated as indicated. Cell viability was analyzed in full or cystine (Cys_2_)-free medium co-treated with 10058-F4 (30 μM, **F3680, Sigma-Aldrich**), z-VAD-fmk (30 μM; sc-3067, Santa Cruz), bafilomycin A1 (200 nM; sc-201550, Santa Cruz), necrostatin-1 (20 μM; N9037, Sigma-Aldrich), ferrostatin-1(fer-1; 5 μM, SML0583, Sigma Aldrich), Trolox (100 μM; 238813; Sigma-Aldrich), or ciclopirox olamine (CPX; 1 μM, sc-204688, Santa Cruz). Cells were fixed with ice-cold 10% trichloroacetic acid (TCA) for 1 h, stained with 0.054 % w/v SRB for 30 min, and absorbance measured at 535 nm using a Tecan Ultra plate reader (Tecan, Maennedorf, Switzerland) at indicated time points. DNA content analysis was performed by fixing cells with 4 % PFA and staining with FxCycle Violet Stain (Thermo Fisher Scientific) followed by FACS using a MACSQuant Flow Cytometer (Miltenyi Biotec, Bergisch Gladbach, Germany).

Doubling times were calculated using two different methods: (i) Impedance measurements were assessed using the RTCA system (Roche) by seeding at different cell densities and registering impedance signals every 20 min. (ii) Standard growth curves were generated by counting cells by FACS (MACSQuant Flow Cytometer) at indicated time points and excluding propidium iodide (PI)-positive cells as necrotic. In both methods, exponential curves were fitted and doubling times calculated.

As another measure of cell viability, CellTiter-Blue cell assays were performed in 96-well plates following the manufacturer’s instructions (Promega, Mannheim, Germany) using a FLUOstar Optima microplate fluorescence reader (BMG LABTECH, Ortenberg, Germany). 24 h after seeding, cells were treated and simultaneously induced with 1 μg/ml doxycycline (Dox) (195044, MP Biomedicals, LLC, Illkirch, France) as appropriate. D9 (synthesized and provided by Bayer Pharma AG, Berlin, Germany), erastin (Cay-17754, Biomol GmbH, Hamburg, Germany), RSL-3 (200 nM; Medchemexpress Europe, Sollentuna, Sweden), SAS (599-79-1, Sigma-Aldrich), DL-Propargylglycine (PPG 1 mM; P7888, Sigma-Aldrich), fer-1 and Fe (VENOFER, 20 mg/ml, Aca Müller/Adag Pharma AG, Gottmadingen, Germany) were used as indicated, and viability determined 72 h after treatment/induction. To assess the effects of Cys_2_ deprivation on cell viability, cells were washed and fed with Cys_2_-free DMEM (Gibco) supplemented with 10 % dialyzed FCS (Gibco), 201.3 μM Met (63-68-3, Sigma-Aldrich) and 4 mM glutamine (Gln; 25030081, Gibco) 48 h after seeding. To determine cell death rescue potential, Cys_2_-deprived cells were also co-treated with homocysteine (Hcy; 69453, Sigma-Aldrich), cystathionine (Cysta; C7505, Sigma-Aldrich) or GSH. We further compared effects on cell viability following Cys_2_ and Gln depletion. Fluorescence was read (540/580nm) 24 h and 48 h after deprivation/treatment.

#### Colony formation assay

IMR5/75-(5,000 cells/well) and SH-EP-AHCYsh (1,000 cells/well) were seeded in six-well plates and simultaneously treated with doxycycline (1 μg/ml) to induce the *AHCY*-targeting shRNA. Cells were fixed (11% glutaraldehyde; Sigma Aldrich) and Giemsa-stained five (SH-EP-AHCYsh) or seven days (IMR5/75-AHCYsh) later. Colony counting was performed using a Gel Doc Documentation System and Quantity One software (Bio-Rad) and quantification using Excel software.

### Metabolite profiling and protein analysis

#### Metabolite Analysis by Gas Chromatography/Mass Spectrometry (GC/MS)

##### Extraction

Approximately 6×10^6^ cells of MYCN-high and MYCN-low expressing IMR5/75, including 10058-F4-(30 μM; F3680; Sigma-Aldrich), or DMSO-treated MYCN-high IMR5/75, were harvested and intracellular GSH and metabolites were measured. Intracellular glutathione levels were further measured as previously described^43^.

Harvested cells were washed twice with 0.9 % ice-cold NaCl solution and snap-frozen in liquid nitrogen. Frozen pellets were extracted in 180 μl of methanol with vigorous shaking for 15 min at 70°C. As internal standard, 5 μl ribitol (0.2 mg/ml, A5502, Sigma Aldrich) were added to each sample. Polar and organic phases were separated with 100 μl chloroform (shaking samples for 5 min at 37°C), and 200 μl water per sample. Following centrifugation (11,000× g, 10 min), 300 μl of the polar (upper) phase were dried in a vacuum concentrator (Eppendorf Concentrator Plus) without heating for derivatization.

##### Derivatization (Methoximation and Silylation)

Pellets were re-dissolved in 20 μl methoximation reagent containing 20 mg/ml methoxyamine hydrochloride (226904, Sigma-Aldrich) in pyridine (270970, Sigma-Aldrich) and incubated for 2 h at 37°C with shaking. For silylation, 32.2 μl *N*-methyl-*N*-(trimethylsilyl)trifluoroacetamide (MSTFA; M7891, Sigma-Aldrich) and 2.8 μl Alkane Standard Mixture (50 mg/ml C_10_ – C_40_; 68281, Fluka) were added to each sample. After incubation for 30 min at 37°C, samples were transferred to glass vials for GC/MS analysis.

##### Gas Chromatography/Mass Spectrometry (GC/MS) Analysis

A GC/MS-QP2010 Plus (Shimadzu^®^) fitted with a Zebron ZB 5MS column (Phenomenex^®^; 30 m × 0.25 mm × 0.25 μm) was used for GC/MS analyses. The GC was operated with an injection temperature of 230°C and 2 μl of each sample were injected with split 10 mode. The GC temperature program started with a 1 min hold at 70°C followed by a 6°C/min ramp to 310°C, a 20°C/min ramp to 330°C and a bake-out for 5 min at 330°C using helium as carrier gas with constant linear velocity. MS was operated with ion source and interface temperatures of 250°C and a scan range (m/z) of 40–1000 with an event time of 0.3 sec. “GCMS solution” software (Shimadzu^®^) was used for data processing.

#### Quantification of Amino Acids

Pellets of 2×10^6^ cells were extracted with 0.1 ml ice-cold 0.1 M HCl. Non-thiol-containing amino acids were quantified after specific fluorescent labeling with AccQ-TagTM (Waters, Eschborn, Germany) as described by Yang et al^44^. Cysteine levels were determined after labeling with monobromobimane (Calbiochem, Merck, Darmstadt, Germany) as described before^45^.

#### Western Blot Analysis of Proteins in Cell Extracts

Whole cell lysates were prepared and protein expression visualized as previously described^46^. Protein lysates (50 μg/lane) were separated on 12.5 % SDSPAGE. Blots were probed with antibodies directed against MYCN (#sc-53993, Santa Cruz, 1:1000), c-MYC (ab32072, Abcam, 1:1000), SAHH (A-11) (AHCY antibody, sc-271389, Santa Cruz, 1:1000) or β-actin-conjugated (ab20272, Abcam, 1:5000). HSR-peroxidase labeled anti-mouse (115-035-003, Dianova, 1:1000) or anti-rabbit (111-035-144, Dianova, 1:1000) antibodies were used as secondary antibodies. Proteins were visualized using ECL detection reagents (Amersham/GE Healthcare, Freiburg, Germany) and a chemiluminescence reader (VILBER, Eberhardzell, Germany). Protein quantification was performed using ImageJ software (https://imagej.net).

### Flow cytometry

#### Analysis of Intracellular ROS Level

Approximately 10^5^ cells were seeded in six-well plates. MYCN-low populations were established by incubating cells with 1 μg/ml doxycycline at least 48 h prior to further treatment. Cells were then fed either with full or cystine-free medium and co-treated with fer-1 (5 μM);, liproxstatin-1 (1 μM; SML1414; Sigma-Aldrich), CPX (1 μM), Trolox (100 μM) or glutathione (GSH; 2 mM, **G4251, Sigma-Aldrich**) for 20 h. Before lipid-ROS being analyzed, medium was removed, C11-BODIPY (581/591) diluted in Hank’s Balanced Salt Solution (HBSS; Gibco) and added to wells at a final concentration of 4 μM. After 15 min staining at 37° C inside the tissue culture incubator, cells were harvested gently and levels of lipid-ROS immediately analyzed using a BD FACS Aria™ III cell sorter. In addition, total intracellular reactive oxygen species (ROS) levels in Cys_2_-deprived cells were determined using CellROX^®^ (Thermo Fisher Scientific). Approximately 10^5^ cells were seeded in 12-well plates. ‘MYCN-low’ populations were established by incubating cells with 1 μg/ml doxycycline, at least 48 h prior to further treatment. To test the effect of MYC inhibition on ROS levels, ‘MYC-high’ cells were treated with 10058-F4 (30 μM for 24 h). Cells were then fed either with full or Cys_2_-free medium 24 h before CellROX^®^ was added (5 μM final concentration). After 30 min of staining in a cell culture incubator, cells were harvested gently and immediately analyzed using a BD FACS Aria™ III cell sorter to measure the signal intensity directly proportional to the level of intracellular ROS.

### MYCN synthetic lethal screen

#### Large-Scale Druggable Genome siRNA Screen

For high-throughput screening, a Silencer Select siRNA custom library (#4404034, Ambion) was used encompassing 31,242 unpooled siRNAs targeting 10,414 genes (3 siRNAs per gene), Lipofectamine RNAiMax transfection reagent (Life Technologies) only and On-TARGETplus Non-targeting siRNA #1 (Dharmacon) served as negative transfection controls, PLK1 (Silencer Select siRNA #1, Ambion) as positive control. Liquid reverse transfection was performed in 384-well plates using a Freedom EVO 200 robotic platform with a 384 MultiChannel Arm, 8-channel Liquid Handling Arm, Robotic Manipulator Arm and EVOware Plus software (TECAN) as follows: Lipofectamine RNAiMax and siRNA were diluted in Opti-MEM (Life Technologies) at appropriate concentrations, combined, and incubated for 15-40 min before adding *MYCN*-amplified IMR5/75 cells stably expressing a doxycycline-regulable MYCN shRNA (2,100 cells/well) using a Multidrop dispenser (Thermo Fisher Scientific). Two treatment conditions were screened in triplicate: (A) culture medium only (IMR5/75 ‘MYCN-high’), and (B) plus doxycycline to induce the shRNA targeting MYCN (IMR5/75 ‘MYCN-low’). After seeding cells into the transfection mixture, medium only (‘MYCN-high’ transfection plates) or doxycycline (1 μg/ml final concentration, ‘MYCN-low’ transfection plates) was added using a Tecan Evo 200 robotic platform. 96 h after transfection, cells were fixed with 11 % glutaraldehyde and subsequently Hoechst-stained (10 mg/ml stock in 1× PBS, 1:2500; Invitrogen) using a Tecan Evo 200 robotic platform. The number of Hoechst-positive cell nuclei was determined using an OPERA fluorescence microscope based on nine sites per well and a BHC in-house program. We applied Redundant siRNA Activity (RSA) to the ratio between MYCN high and MYCN low data to select top ranked hits^47^ and a false discovery rate of (FDR) of 0.2. Raw and processed data of the druggable whole-genome screen will be made available after publication.

### Transcript profiling tumors and cell lines

#### RNA-sequencing (RNA-seq)

Total RNA was isolated using the miRNeasy Mini Kit (Qiagen) and depleted from ribosomal RNAs using the Ribo-Zero rRNA Removal Kit (Illumina) according to the manufacturers’ protocols. RNA libraries were prepared using the NEBNext Ultra Directional RNA Library Prep Kit for Illumina (New England BioLabs) with following changes to the manufacturer’s instructions: RNA was fragmented for 20 min at 94°C, followed by first strand cDNA synthesis for 10 min at 25°C, 50 min at 42°C and 15 min at 70°C. Adapter-ligated DNA was size-selected with a bead:DNA ratio of 0.4 (AMPure XP beads, Beckman Coulter), removing index primer and short fragments. Quality, quantity and sizing (approximately 320 bp) of the RNA library were checked using a DNA High Sensitivity DNA chip run on a 2100 Bioanalyzer sequencing platform (German Cancer Research Center Core facility).

#### Data analysis RNA-seq

RNA-seq expression profiles from 498 primary neuroblastomas^48^ (GSE62564) were analyzed. The Wilcoxon rank sum test was used to test an association between candidate gene expression and amplified *MYCN* oncogene. Maximally selected log-rank statistics were used to describe the relationship between patient survival and candidate gene expression, and resulting expression cut points were used for dichotomization. Survival curves were estimated using the Kaplan-Meier method. Cox regression was used to investigate the prognostic power of candidate gene expression adjusting for established prognostic variables. Parameter estimate shrinkage was applied to correct for potential overestimation of the hazard ratio estimate due to cut point selection^49^. Pearson correlation coefficients were calculated to estimate linear dependence between expression values of candidate genes.

Raw RNA-seq sequences neuroblastoma cell lines and model systems were mapped to the UCSC hg19 genome by STAR^50^ with default parameters. The corresponding bigwig files of each sample were calculated by RSeQC^51^ and normalized to the wigsum of 100000000. RNA-seq sequences from the IMR5/75 MYCN-high and MYCN-low cells were processed as previously described^32^.

### Epigenetic characterization of tumors and cell lines

#### Chromatin immunoprecipitation DNA-sequencing (ChIP-seq) analysis of histone modifications in neuroblastoma primary tumors and cell lines

Formaldehyde cross-linking of cells, cell lysis, sonication, chromatin immunoprecipitation (IP) and library preparation were performed as described previously^52^, starting with approximately 4×10^6^ cells (1×10^6^ cells per individual IP). Direct cell lysis for each sample was achieved by 30 min incubation on ice in 950 μL RIPA I (10 mM Tris-HCl pH 8.0, 1 mM EDTA pH 8.0, 140 mM NaCl, 0.2 % SDS, 0.1 % DOC). Tissue disruption, formaldehyde fixation and sonication of tumor material was done as previously published^53^, using approximately 30 mg of fresh-frozen tumor tissue per individual ChIP-seq experiment. All subsequent steps were performed analogous to cell line experiments. A Bioruptor Plus sonication device (Diagenode) was used for high intensity sonication for 30-60 min each with 30s on and 30s off intervals. Following antibodies were used for IP: H3K4me3 (ab8580, Abcam), H3K4me1 (ab8895, Abcam), H3K27Ac (ab4729, Abcam), H3K27me3 (39155, Active Motif), H3K36me3 (ab9050, Abcam), H3K9me3 (ab8898, Abcam). Library preparation was performed using the NEBNext Ultra DNA Library Prep Kit (New England Biolabs) according to the manufacturer’s protocol. Samples were mixed in equal molar ratios and sequenced on an Illumina sequencing platform.

#### ChIPmentation of MYCN transcription factor in neuroblastoma cell lines

Formaldehyde cross-linking, cell lysis, sonication and chromatin immunoprecipitation were performed as previously described^54^, adding the Chipmentation module by Schmidl et al.^55^ with following changes: a Bioruptor Plus with automated cooling (4°C) was used for high intensity sonication (20-30 min each with 30 sec on and 30 sec off intervals), and 10 μg MYCN antibody (MYCN (sc-53993, Santa Cruz) and 10^6^ cells for chromatin immunoprecipitation. The tagmentation reaction (Illumina Nextera DNA library Prep Kit) was performed at 37°C for 1 min with the bead-bound chromatin sample or 5 ng purified input DNA for normalization. After de-crosslinking, purified samples were amplified using dual index barcodes of the Nextera Index kit and 13 PCR cycles. Enriched libraries were purified and pooled. ChIPmentation libraries were sequenced (50 bases single-end) on the Illumina sequencing platform (German Cancer Research Center Core facility).

#### Data analysis ChIP-seq and ChIPmentation

Single end reads were aligned to the hg19 genome using Bowtie2 (version 2.1.0), keeping uniquely aligned reads only. BAM-Files of aligned reads were further processed using the deepTools suite^56^. Input files were subtracted from treatment files using the bamCompare tool, applying the SES method for normalization of signal to noise. Resulting signals were normalized to an average 1X coverage to produce signal (bigWig) files. Peaks were called using the MACS 1.4 tool using default parameters. Data sets from cell lines and primary tumors will be made available after publication.

#### DNA methylation analysis

DNA methylation and gene expression data from 105 primary neuroblastomas assessed by Infinium HumanMethylation450 BeadChips and 44k Agilent oligonucleotide microarrays^13^ (GEO accession GSE73518) were analyzed for candidate loci. For investigating correlation between DNA methylation and expression, maximally selected Wilcoxon rank-sum statistics were used estimating CpG methylation cut points that separate patient groups with differential gene expression. To test the null hypothesis that odds ratios estimating the association of methylation-separated patient subgroups with high-risk disease were 1, a Fisher exact test was used. R2 Genomics Analysis and Visualization Platform (http://r2.amc.nl) was used to visualize expression/methylation of selected gene/CpG pairs.

### Tumor proteome analysis

Patient samples were lysed with 2 % SDS, 50mM ammonium bicarbonate buffer and complete, mini, EDTA-free Protease Inhibitor Cocktail. Samples were homogenized at RT 15 sec for 7 cycles with a 5 sec pause using FastPrep-24™ 5G Homogenizer and heated to 95° C for 5 min. Freeze-thaw cycles were repeated seven times. Benzonase was added to each sample and the lysate was clarified by centrifugation at 16 rcf for 30 min at 4°C. Protein concentration was then measured using the Bio-Rad DC Protein assay. From each sample, 50 μg of protein were taken in duplicate. Proteins were reduced with 10 mM DTT for 30 min and alkylated with 55 mM iodoacetamide for 30 min. Wessel-Flügge precipitation was performed as previously described^57^. The retrieved protein pellet was resuspended in 6 M Urea, 2 M Thiourea in 10 mM HEPES (pH 8). Proteins were then digested with LysC for 2 h; samples were diluted in 50mM ammonium bicarbonate buffer to reach less than 2M urea before trypsin was added for overnight digestion. The peptide solution was acidified and fractionated by SCX on StageTips as described previously^58^. Samples were then separated by reversed phase HPLC on a 2,000 mm monolithic column and analyzed online by high resolution mass spectrometry on a Q Exactive plus mass instrument as described previously^59^. The resulting raw files were analyzed using MaxQuant software version 1.5.5.1^60^. Default settings were used with ‘match between runs’ function. Data was searched against a Human Uniprot database (2014-10) and common contaminants at a false discovery rate of 1 % at both the peptide and protein level. The resulting text files were filtered to exclude reverse database hits, potential contaminants, and proteins only identified by site. Plotting and statistics were done using Perseus software version 1.5.5.3.^61^.

